# Bursts of rapid diversification, dispersals out of southern Africa, and two origins of dioecy punctuate the evolution of *Asparagus*

**DOI:** 10.1101/2024.07.25.605174

**Authors:** Philip C. Bentz, John E. Burrows, Sandra M. Burrows, Eshchar Mizrachi, Zhengjie Liu, Jun-Bo Yang, Zichao Mao, Margot Popecki, Ole Seberg, Gitte Petersen, Jim Leebens-Mack

## Abstract

The genus *Asparagus* arose approximately 9–15 million years ago (Ma) and transitions from hermaphroditism to dioecy (separate sexes) occurred ∼3–4 Ma. Roughly 27% of extant *Asparagus* species are dioecious, while the remaining are bisexual with monoclinous flowers. As such, *Asparagus* is an ideal model taxon for studying early stages of dioecy and sex chromosome evolution in plants. Until now, however, understanding of diversification and shifts from hermaphroditism to dioecy in *Asparagus* has been hampered by the lack of robust species tree estimates for the genus. In this study, a genus-wide phylogenomic analysis including 1726 nuclear loci and comprehensive species sampling supports two independent origins of dioecy in *Asparagus*—first in a widely distributed Eurasian clade, then again in a clade restricted to the Mediterranean Basin. Modeling of ancestral biogeography indicates that both dioecy origins were associated with range expansion out of southern Africa. Our findings also revealed several bursts of diversification across the phylogeny, including an initial radiation in southern Africa that gave rise to 12 major clades in the genus, and more recent radiations that have resulted in paraphyly and polyphyly among closely related species, as expected given active speciation processes. Lastly, we report that the geographic origin of domesticated garden asparagus (*Asparagus officinalis* L.) was likely in western Asia near the Mediterranean Sea. The presented phylogenomic framework for *Asparagus* is foundational for ongoing genomic investigations of diversification and functional trait evolution in the genus and contributes to its utility for understanding the origin and early evolution of dioecy and sex chromosomes.

**Significance Statement:** *Asparagus* is an important model system for studying dioecy (separate sexes) evolution in plants. *Asparagus* taxonomy has been challenging, likely due to rapid species diversifications leading to highly variable species with complicated relationships that are impossible to resolve with limited DNA-sequence data. Using phylogenomics and the largest species sampling to date, we show that all *Asparagus* lineages originated from an initial radiation in southern Africa and that separate range expansions out of southern Africa set the stage for two distinct origins of dioecy in *Asparagus*. Our findings provide a deeper understanding of species diversification and the role of long-distance dispersals in the evolution of dioecy. This study also illustrates the utility of phylogenomics for elucidating past and present speciation processes.

## Introduction

The genus *Asparagus* Tourn. ex L. encompasses over 215 species and can be found in many different habitats spanning nearly all of Africa and Eurasia (1). The most well-known species in the genus is garden asparagus (*Asparagus officinalis* L.) which is an important vegetable crop cultivated across the globe (2) and a model system for studying dioecy and sex chromosome evolution in angiosperms (3–10). *Asparagus* is in the Asparagoideae subfamily of Asparagaceae, along with one other genus— *Hemiphylacus* S. Watson (11). The biodiversity hotspot for *Asparagus* is southern Africa but is distributed across the eastern hemisphere (12), whereas its sister genus, *Hemiphylacus*, is endemic to Mexico (13). Approximately 73% of *Asparagus* species— and all five *Hemiphylacus* species—are bisexual with monoclinous flowers, while the rest of *Asparagus* are dioecious (14). Previous studies have failed to confidently resolve species relationships across the *Asparagus* phylogeny despite many attempts using few organellar and/or nuclear loci (12, 15–19). Ancestral rapid radiations likely account for the difficulty of phylogenetic inference in *Asparagus* due to incomplete lineage sorting (ILS) (12, 14). These phylogenetic challenges are compounded by the young origins of the genus (i.e., ∼9–15 Ma) and low substitution rate for the plastome (14). Poor resolution of species relationships in *Asparagus* has stymied robust inference for the origin(s) of dioecy in the genus, with some analyses supporting a single origin (16, 18) and others opening the possibility for two separate origins (12, 14).

To explore evolutionary relationships and ancestral biogeography across the genus *Asparagus*, we employed a Hyb-Seq approach with Asparagaceae1726—a probe set targeting 1726 nuclear loci conserved in low copy numbers across Asparagaceae (52)—and the most comprehensive species sampling of *Asparagus* to date. Hyb-Seq, or target sequence capture, is a cost-effective, reduced representation sequencing strategy that utilizes RNA probes designed to select and enrich for specific genes of interest (20). In this study, we i) investigate how rapid radiations and range expansion out of southern Africa may have contributed to the origin and diversification of major *Asparagus* lineages; ii) explore possible explanations for complicated speciation patterns and taxonomic uncertainties; iii) test the hypothesized multiple origins of dioecy in *Asparagus* (12, 14); and iv) infer the geographic origin of domesticated garden asparagus.

## Results

### Species sampling and target sequence capture

In this study, we assembled the most comprehensive species sampling to date for phylogenomic investigation of *Asparagus*, including 314 accessions representing 158 *Asparagus* species (Table 1). Our sampling effort yielded phylogenomic support for 28 new species (+2 subsp.), the resurrection of nine previously described species, and elevation of three varieties to the species level (Supporting Information, Appendix S1). Formal taxonomic revision is outside the scope of the current study but these new southern African *Asparagus* species, as well as other taxonomic updates, are described in a forthcoming publication (J.E. Burrows & S.M. Burrows, in prep.). After performing Hyb-Seq with the Asparagaceae1726 bait set (52), we recovered >1300 orthologous target loci in all samples (mean=1680) (Appendix S2). A small fraction target loci (0.9– 2.8%) exhibited gene copy number variation (i.e., paralogs) among taxa (Results S1; Appendix S2). However, analyses with and without paralogs resulted in congruent topologies and branch support, suggesting limited effect on phylogenomic inference in *Asparagus* (52).

**Table 1.** Species sampling across 12 major clades in *Asparagus* and outgroups.

| Clade | Samples | New species | Total species |
| --- | --- | --- | --- |
| ***Asparagus* clades** |  |  |  |
| Africani | 33 | 1 | 13 (+1 subsp.) |
| Asparagus | 69 | 1 | 32 |
| Exuviali | 6 | 1 | 3 |
| Mackowanii | 2 | 0 | 2 |
| Myrsiphyllum | 33 | 4 | 12 |
| Nutlet | 5 | 1 | 4 |
| Racemose | 90 | 17 | 49 (+1 var.) |
| Retrofracti | 16 | 1 | 8 (+1 subsp.) |
| Scandens | 5 | 0 | 2 |
| Setaceus | 19 | 1 (+1 subsp.) | 9 (+1 subsp.) |
| Suaveolens | 29 | 6 (+1 subsp.) | 16 (+1 subsp.) |
| Sympodioidi | 7 | 0 | 5 |
| *Hemiphylacus* | 1 | 0 | 1 |
| Brodiaeoideae (outgroup) | 1 | 0 | 1 |
| Scilloideae (outgroup) | 1 | 0 | 1 |
| Lomandroideae (outgroup) | 1 | 0 | 1 |
| **Total** | **318** | **33 (+2 subsp.)** | **162 (+4 subsp.)** |

**Note:** Outgroup species include *Hemiphylacus* – the sister genus to *Asparagus* and sole other member of the Asparagaceae subfamily Asparagoideae – and samples from three different Asparagaceae subfamilies. New species circumscriptions are presented in a forthcoming monograph and revision of southern African *Asparagus* (J.E. Burrows & S.M. Burrows, in prep.).

### The *Asparagus* phylogeny

Phylogenomic analyses yielded strong support (i.e., local posterior probability [LPP]=1.0) for the earliest split in the *Asparagus* phylogeny, spawning one largely African branch leading to ten major clades and another spawning the Asparagus and Exuviali sister clades (Fig. 1). Most species in the Asparagus clade are dioecious and are widespread across Eurasia and the Mediterranean Basin, whereas all other major clades are solely bisexual and largely restricted to Africa. Across the species tree, several paraphyletic species were revealed (Fig. S1; Results S1) some of which exhibit signatures of an ongoing budding speciation process (see Discussion). Multiple polyphyletic species were also observed the *Asparagus* phylogeny, including a few highly polymorphic species known to be taxonomically troublesome (i.e., *A. asparagoides*, *A. densiflorus* and *A. setaceus*) (Results S1).

**Figure 1.**
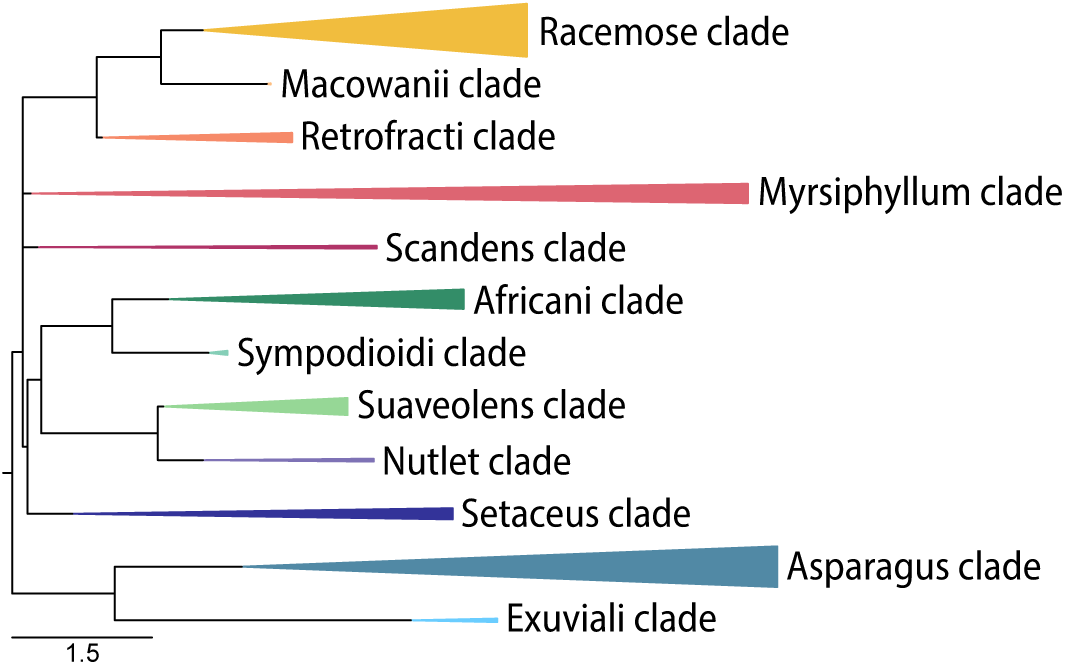
Relationships among 12 major clades representing nearly the full extent of species diversity and geographic range of the genus *Asparagus*. All clades are solely composed of bisexual species exhibiting monoclinous flowers, except for the Asparagus clade in which dioecy evolved twice independently.Summary tree based on larger phylogeny of 318 accessions (162 species) inferred from 1726 nuclear genes (52) using wASTRAL-unweighted v1.16.3.4 (57) by optimizing the objective function of ASTRAL-III v.5.7.8 (58). Triangular branch widths were scaled according to number of samples and initiation of tapering represents crown group radiations. Branch lengths/scale bar correspond to coalescent time units. Bifurcations were collapsed according to a polytomy test with ASTRAL-III (*p*-value >0.05). All illustrated branches had local posterior probability (LPP) support ≥0.99, except when including *A. fasciculatus* (sister species to the rest of Myrsiphyllum) in Myrsiphyllum (LPP=0.64), otherwise LPP=0.99 for the clade.

Across the final species tree (Fig. S1), gene tree discordance (Fig. S2) was relatively low (ASTRAL normalized quartet score=0.77), and quartets showed limited signs of deviation from assumptions of the multispecies coalescent, which accounts for ILS between speciation events (71, 72, 73). For example, ancestral gene flow between diverging lineages can result in low Quartet Differential (QD) scores (47) (QD=0 when all discordant quartets at a node support one of the two alternative resolutions and QD=1 when quartet frequencies for both alternatives are equal). The average QD across all nodes in the *Asparagus* phylogeny was 0.87 (Fig. S2). Furthermore, many of the nodes exhibiting QD<0.50 involved bifurcations between samples of the same species (Fig. S2). At the same time, the *Asparagus* phylogeny included 27 nodes for which we were unable to reject a polytomy (i.e., nodes with more than two daughter lineages) (Fig. S3). In a fully bifurcating tree, branches around these nodes exhibited low LPP and short branch lengths (Fig. S4), and similar gene tree quartet support for both alternative resolutions, rather than skewed support favoring one alternative topology over the other (Figs. S2, S5).

A polytomy could not be rejected (*p-*value=0.07: Fig. S3) among the Scandens, Myrsiphyllum, and Racemose-Macowanii-Retrofracti clades; due to low support (LPP=0.83) and short branch length subtending a Myrsiphyllum-Scandens clade (Fig. S4), which also exhibited similar quartet support for alternative resolutions (QD=0.94) (Figs. S2, S5). Additionally, a polytomy could not be rejected (*p-*value=0.27) along the African *Asparagus* backbone—or branch subtending a i) Africani-Sympodioidi-Suaveolens-Nutlet-Setaceus clade; ii) Myrsiphyllum-Scandens clade; and iii) Racemose-Macowanii-Retrofracti clade—due to low bifurcating support between the Myrsiphyllum-Scandens and Racemose-Macowanii-Retrofracti clades (LPP=0.82; QD=0.98) (Figs. S2–S5). Collapsing of the two branches described above resulted in a polytomy with four leaves forming the backbone of the main African, bisexual clade in the genus (Fig. 1; Fig. S1). Aside from Myrsiphyllum, all other major clades were strongly supported (LPP≥0.99) (Fig. S1). The inclusion of *A. fasciculatus* (a lineage sister to the rest of Myrsiphyllum) in the Myrsiphyllum clade was weakly supported (LPP=0.64) but monophyly of the remainder of Myrsiphyllum was strongly supported (LPP=0.99) (Fig. S1).

The Asparagus clade was strongly supported and showed virtually no skew in alternative quartet support (QD=0.94) (Fig. S2). Within the Asparagus clade lies three smaller, well-supported (LPP=1.0) clades, including a clade of four bisexual species that are distributed across central-southern Africa (Flagellaris clade: Fig. 2); and two entirely dioecious clades including a broadly distributed Eurasian clade, and a less speciose and more narrowly distributed Mediterranean Basin clade (Fig. 2). All three of those nested clades were also supported by most quartets with limited to no skew between alternative resolutions (Figs. S2, S5). The Medit. Basin dioecy clade and Flagellaris bisexual clade together form a clade that is sister to the Eurasian dioecy clade (LPP=1.0; QD=0.98) (Figs. 2, S2). Lastly, the Exuviali clade, composed of three species restricted to southern Africa (Fig. 2), was highly supported (LPP=1.0; QD=1.0), and placed sister to the Asparagus clade (Figs. 1, S2).

**Figure 2.**
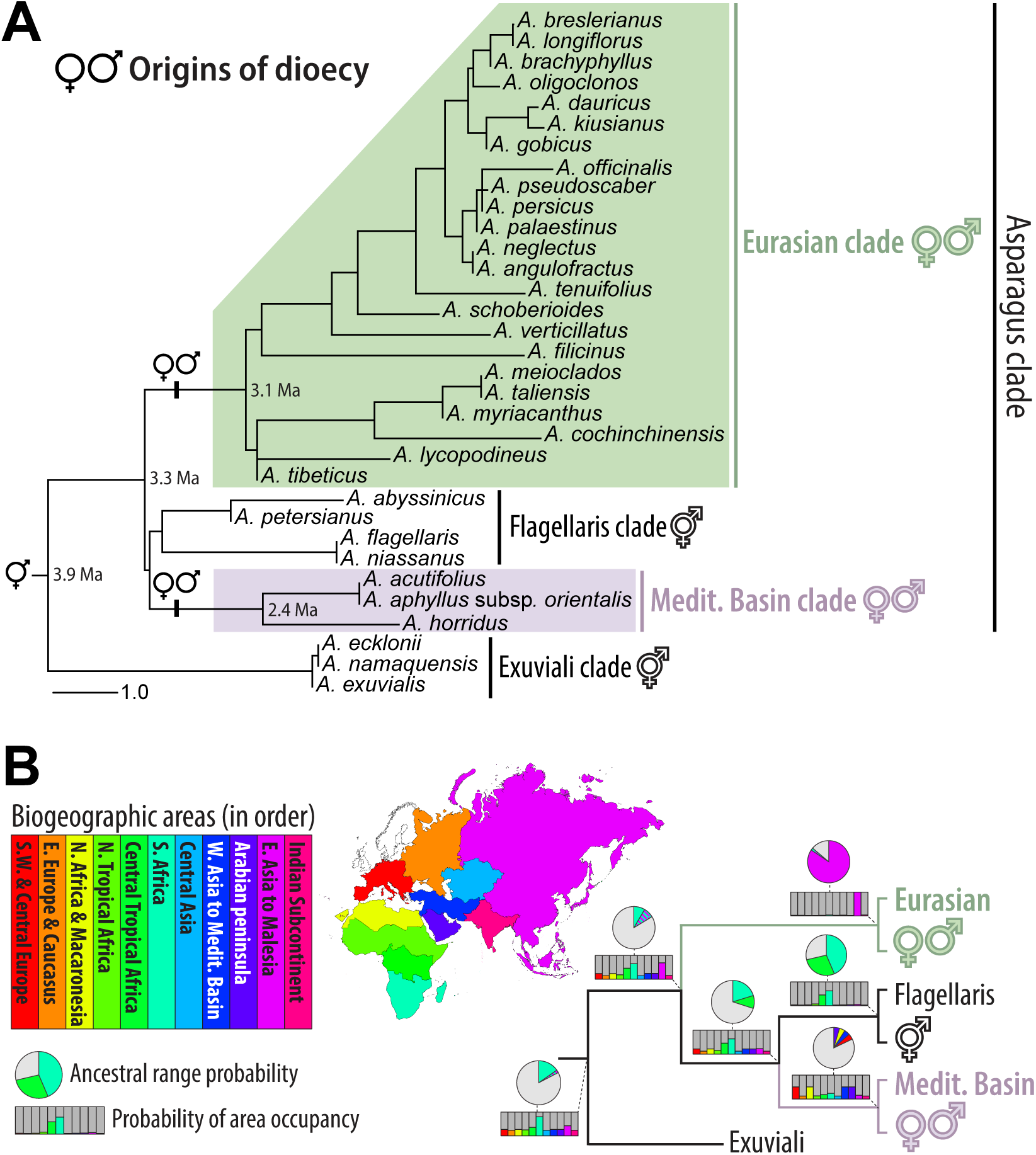
Phylogeny of the Asparagus clade **(A)** and ancestral range estimation **(B)** showing support for two independent origins of dioecy in the genus *Asparagus*. **A)** Species tree supports two separate dioecious clades nested in the Asparagus clade: a Eurasian clade (green) and a Mediterranean Basin clade (purple). Local posterior probabilities equaled 1.0 for all branches. When present, multiple samples for a species were collapsed into a single terminal branch. Bifurcations were collapsed according to a polytomy test with ASTRAL-III v.5.7.8 (58). Divergence time estimates (Ma=million years ago) are plotted at concordant nodes from Bentz et al. (14) and branch lengths/scale bar represent coalescent time units. **B)** Ancestral range estimation suggests that the stem ancestor of the Medit. Basin dioecy clade dispersed out of southern or central Africa independently of the Eurasian dioecy clade. Cladogram shows summarized results from BioGeoBEARS v.1.1.1 (61, 62) plotted at relevant nodes. Pies show the top two ancestral *range* probabilities in color (grey=all other possibilities) for internal nodes, except at the crown node of the Medit. Basin dioecy clade, which had four nearly equal top probabilities (all ∼5%). Bar charts show the relative probability of geographic *area* occupancy (i.e., probability that a *range* included any of the 11 predefined *areas*). *Area* = discrete geographic region. *Range* = species distribution encompassing any combination of *areas.*

### Ancestral biogeography estimation

The best fit biogeographic model for this dataset was Dispersal–Extinction–Cladogenesis (DEC) (Table S1) and allowing for founder events within clades (parameter *j*) significantly improved model fit relative to the DEC model (*p*-value=0.026) (Table S2). Ancestral range inference under the DEC+J model implicates southern Africa as the most probable center of origin for all 12 major clades in *Asparagus* (Fig. S6a; Appendix S3). Our results indicate that at least eight independent range shifts, across the genus, resulted in dispersal out of Africa (Fig. S6a; Appendix S3). One of these shifts out of Africa occurred following the split between the Exuviali and Asparagus clades, in the Asparagus clade crown group, which most likely originated in southern Africa (Fig. 2; Table 2). However, a disjunct range including southern Africa and eastern Asia was almost as probable for the crown group of the Asparagus clade (Figs. 2, S6b; Table 2), which is practically impossible considering the geographic distance between these regions. Such uncertainty at the crown of the Asparagus clade may be explained by sampling bias (i.e., eastern Asia is the biodiversity hotspot for the Eurasian clade and is more speciose than its sister clade) and can be further assessed at surrounding splits in the tree, which suggest a transition from Africa to Asia occurred following cladogenesis of the Eurasian lineage (Figs. S6, S7a; Appendix S3). Nonetheless, the ancestral range for the Exuviali-Asparagus crown group was inferred as southern Africa, whereas the crown range for the Eurasian dioecy clade was placed in eastern Asia with high confidence (Fig. 2; Table 2). Also in the Asparagus clade, a separate ancestral range shift was estimated at the stem branch of the Medit. Basin dioecy clade, initiating out of either southern or central Africa and ending with colonization of the Arabian Peninsula or Medit. Basin by the crown group (Figs. 2, S6–S7; Table 2).

**Table 2.** Ancestral range estimations for nodes implicated in the two origins of dioecy in *Asparagus* and origin of garden asparagus (*Asparagus officinalis* L.).

|  | Asparagus<br>genus crown | Asparagus<br>clade stem | Asparagus<br>clade crown | Eurasian<br>clade crown | garden<br>asparagus<br>clade stem<sup>(a)</sup> | Medit. Basin<br>clade stem | Medit. Basin<br>clade crown | Flagellaris<br>clade crown |
| --- | --- | --- | --- | --- | --- | --- | --- | --- |
| **A) Relative probability of occupancy per geographic area** |  |  |  |  |  |  |  |  |
| **Areas** |  |  |  |  |  |  |  |  |
| S. & Central Europe (D) | 31% | 27% | 24% | 3% | 33% | 21% | **50%** | 4% |
| E. Europe & Caucasus (E) | 29% | 24% | 16% | 3% | 33% | 12% | 13% | 3% |
| N. Africa & Canary Isl. (F) | 31% | 27% | 24% | 2% | 0% | 21% | **50%** | 4% |
| N. Tropical Africa (J) | 30% | 25% | 19% | 2% | 0% | 16% | 13% | 8% |
| Central Tropical Africa (K) | 36% | 36% | **46%** | 3% | 0% | **46%** | 18% | **45%** |
| Central Asia (L) | 29% | 24% | 16% | 3% | 37% | 12% | 13% | 3% |
| S. Africa (M) | **100%** | **80%** | **65%** | 4% | 0% | **64%** | 23% | **62%** |
| W. Asia to Medit. Basin (R) | 31% | 27% | 24% | 3% | **100%** | 21% | **50%** | 4% |
| Arabian Peninsula (T) | 31% | 27% | 24% | 2% | 0% | 22% | **50%** | 4% |
| E. Asia to Malesia (U) | 39% | 42% | **69%** | **100%** | 46% | 23% | 15% | 6% |
| Indian subcontinent (X) | 29% | 24% | 16% | 4% | 37% | 12% | 13% | 3% |
| **B) Most probable ancestral ranges (in order of highest probability)** |  |  |  |  |  |  |  |  |
| **#1** | M | M | M | U | R | M | T | M |
| probability | 9% | 15% | 8% | 84% | 24% | 20% | 5% | 44% |
| **#2** | M+U | M+U | M+U | M+U | R+U | K | R | K |
| probability | 1% | 2% | 8% | 2% | 7% | 10% | 5% | 28% |
| **#3** | K+M | K+M | K+U | U+X | D+<sup>(b)</sup> | M+U | D | K+M |
| probability | 1% | 1% | 4% | 1% | 6% | 2% | 5% | 5% |
| **#4** | M+T | M+T | K | R+U | L+R | K+M | F | J |
| probability | 1% | 1% | 2% | 1% | 4% | 1% | 5% | 1% |

**Note:** **A)** is the relative probability of occupancy for a given node in each of the 11 geographic areas (states) defined for biogeography analysis and illustrated as bar charts in Fig. 2.
<sup>a</sup> Well-supported branch with polytomy among *A. officinalis*, *A. persicus*, *A. pseudoscaberr* (Fig. 2).
<sup>b</sup> E+L+R+U+X.

### Discussion Rapid radiations and dispersals out of Africa

Biogeographic analyses strongly support southern Africa as the ancestral center of origin for the genus *Asparagus* (Table 2), which agrees with previous findings (12). The short branches and polytomous backbone of the *Asparagus* phylogeny indicates that early and rapid diversification in southern Africa gave rise to all extant major clades in the genus (Fig. 1). Following its origin in southern Africa, eight independent range expansion events, in four major clades, led to colonization of Eurasia Fig. S6). One of these expansions out of southern Africa initiated at the crown node of the Asparagus clade and led to a founding event in eastern Asia, where dioecy evolved in the most recent common ancestor (MRCA) of the Eurasian dioecy clade (Fig. 2; Table 2) and was followed by rapid diversification of dioecious lineages and secondary dispersals across the continent (Fig. 3; Results S1). Our results suggest that the MRCA of the Medit. Basin dioecy clade dispersed out of southern Africa after that of the Eurasian dioecy clade, perhaps through the Arabian Peninsula (Figs. 2–3; Table 2).

**Figure 3.**
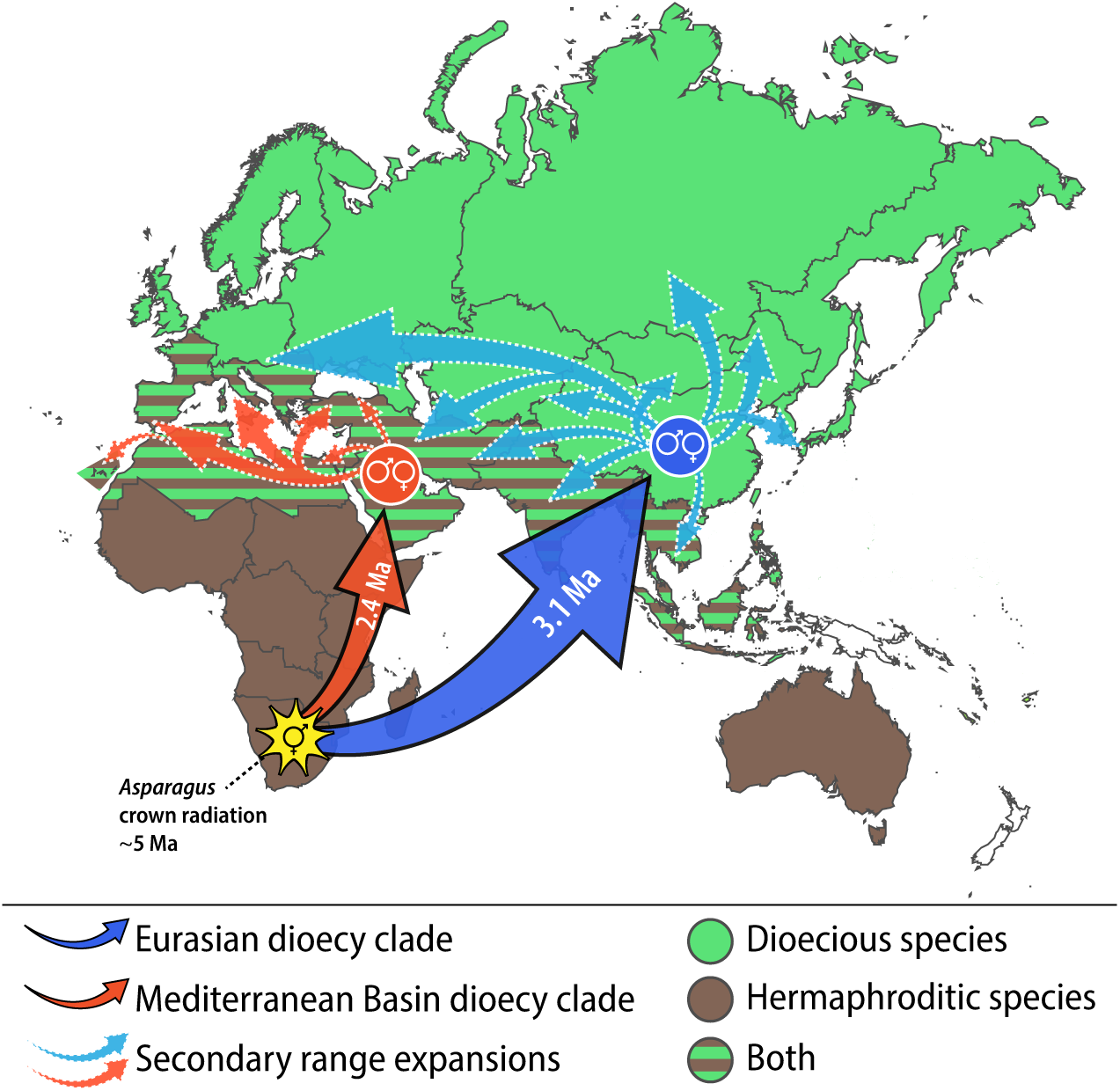
Hypothesized long-distance dispersals out of southern Africa and into Eurasia by separate bisexual ancestral lineages (solid outline arrows), leading to independent founding events and origins of dioecy in two clades of *Asparagus* (red and blue circles). Arrows do not represent specific dispersal routes, which remain uncertain, and are intended to simply point to destinations. The inferred founding events in Eurasia (circles) are based on the most likely ancestral ranges for each dioecious clade. It is nearly as probable that founders of the Medit. Basin dioecy clade first occurred elsewhere in the Medit. Basin, rather than the Arabian Peninsula (Table 2). Divergence time estimates plotted on the larger arrows correspond to mean crown group ages for each dioecy clade from Bentz et al. (14) (Ma=million years ago).

Interestingly, our results suggest that ancestors of extant, non-African bisexual *Asparagus* lineages expanded northeastward out of southern Africa, through the Arabian Peninsula, into Eurasia and that dispersal to Macaronesia occurred independently three times (Results S1). Altogether, our biogeographic analysis suggests that range expansions out of southern Africa occurred independently in various *Asparagus* lineages following an initial radiation that formed all extant major clades (Fig. 1) and alongside clade-specific secondary radiations. Across the *Asparagus* phylogeny, we found virtually no evidence of secondary evolutionary histories (e.g., ancient hybridizations, lineage-specific rate heterogeneity or heterogeneous base compositions, etc.) as indicated by a skew in quartet support for a single alternative resolution, or a low QD (47). This includes the four-way polytomy representing the backbone of the main 10 African clades in the genus (Fig. 1), which implicates only ILS as the cause of gene tree discordance for those nodes.

### Two independent origins of dioecy in *Asparagus*

The inferred *Asparagus* phylogeny resulted in strong support for a bifurcation between the Eurasian dioecy clade and two sister clades: a dioecious Medit. Basin clade and a bisexual clade primarily composed of African species (Flagellaris clade) (Fig. 2). Support for two dioecious clades in *Asparagus* was previously found based on whole plastome sequences, though a polytomy among those and the Flagellaris clade showed higher support compared to bifurcations (14). In contrast, our analysis of 1726 nuclear loci and many more species enabled rejection of a polytomy among those three clades (Fig. 2). Norup et al. (12) also reported two dioecious lineages within the Asparagus clade, but with very poor support. Parsimony mapping of trait shifts across the *Asparagus* phylogeny may imply equal support for a single origin of dioecy—on the stem branch of the Asparagus clade—followed by a loss in the bisexual Flagellaris clade, or two independent origins in the Eurasian and Medit. Basin dioecy clades (Fig. 2). However, when considering extant and ancestral species ranges in the genus, as well as the sister relationship between the Medit. Basin dioecy clade and bisexual Flagellaris clade (Fig. 2), the inference of two separate dioecy origins is strengthened. Biogeographic analysis suggests that if dioecy evolved only once in *Asparagus*, then this would have occurred in Sub-Saharan Africa (see Asparagus clade crown in Fig. 2b). However, the practically complete absence of dioecious *Asparagus* in Africa (outside of the Medit. Basin) suggests that a Sub-Saharan dioecy origin is unlikely.

Ancestral range estimations implicated eight independent range shifts out of southern Africa, two of which are associated with long-distance dispersal events and the evolution of dioecy in the Asparagus clade. Transitions from hermaphroditism to dioecy, following long-distance dispersals and associated with colonization bottlenecks, supports the idea that dioecy may evolve in response to selective pressure to avoid inbreeding and deleterious mutational loads in founding populations (34). A founding population that formed after a long-distance dispersal event may be at increased risk to experience inbreeding depression due to a higher propensity to self, which is thought to be a required condition for long-distance dispersal and colonization of new habitats (37). To that end, long-distance dispersals and founder population bottlenecks are hypothesized to have contributed to the evolution of dioecy in many angiosperms endemic to oceanic islands (38, 39). In *A. officinalis*, some genotypically male (XY) plants bear a combination of strictly male flowers and bisexual flowers (i.e., andromonoecy) that readily self-pollinate and set fruit with no obvious barriers present, and the resulting progeny typically exhibit traits thought to correlate with inbreeding depression (e.g., loss of germinability and vigor) (35, 36). Similarly, andromonoecy and selfing in bisexual flowers has been observed in several other dioecious *Asparagus* (e.g., *A. horridus*, *Asparagus cochinchinensis* [Lour.] Merr., and *Asparagus taliensis* F.T.Wang & Tang ex S.C.Chen). Such high selfing rates in andromonoecious individuals of *A. officinalis* and others may be reminiscent of a bisexual ancestral state with a predisposition to self (35). Therefore, if the ancestrally bisexual founder populations, of extant dioecious lineages of *Asparagus*, exhibited high selfing rates, then inbreeding depression could have promoted selection for outcrossing and the evolution of dioecy in both dioecious clades.

Genomic comparisons between *A. officinalis* (Eurasian dioecy clade) and *Asparagus horridus* L. (Medit. Basin dioecy clade) also support independent origins of dioecy in these clades, since each exhibit novel sex chromosomes that separately evolved from different autosomes (32; Bentz et al., in prep.). Furthermore, no homologous genes exist between the *A. officinalis* and *A. horridus* Y-linked sex-determining regions (SDRs), where the master sex-determining genes reside (e.g., *SOFF* and *aspTDF1* in *A. officinalis*) (5, 32). Expression assays of the male promoter gene, *aspTDF1*, also support the absence of a Y-linked *aspTDF1* in *A. horridus* and another member of the Medit. Basin dioecy clade (*Asparagus acutifolius* L.) as evident by its equal expression in both sexes of each species (33). The presence of completely different sex chromosomes and disparate genetic pathways for sex-determination, between the Eurasian and Medit. Basin dioecy clades, along with the biogeographic and phylogenomic support presented here (Fig. 2; Table 2), suggests strongly that dioecy arose twice independently in *Asparagus* and in association with separate range expansions from southern Africa to Eurasia (Figs. 2–3).

### Active speciation in *Asparagus*

The *Asparagus* phylogeny includes some paraphyletic and polyphyletic species, typically involving narrowly distributed species nested within geographically widespread and highly polymorphic species (Results S1). We identified several examples of paraphyletic species (e.g., Fig. 4) that may be explained as a consequence of species with restricted ranges or ecological niches forming from within a more geographically widespread and phylogenetically diverse (now paraphyletic) extant species. This process of speciation, through genetic divergence of a population from an extant parent species, in association with reproductive isolation, is commonly referred to as “budding speciation” (21).

**Figure 4.**
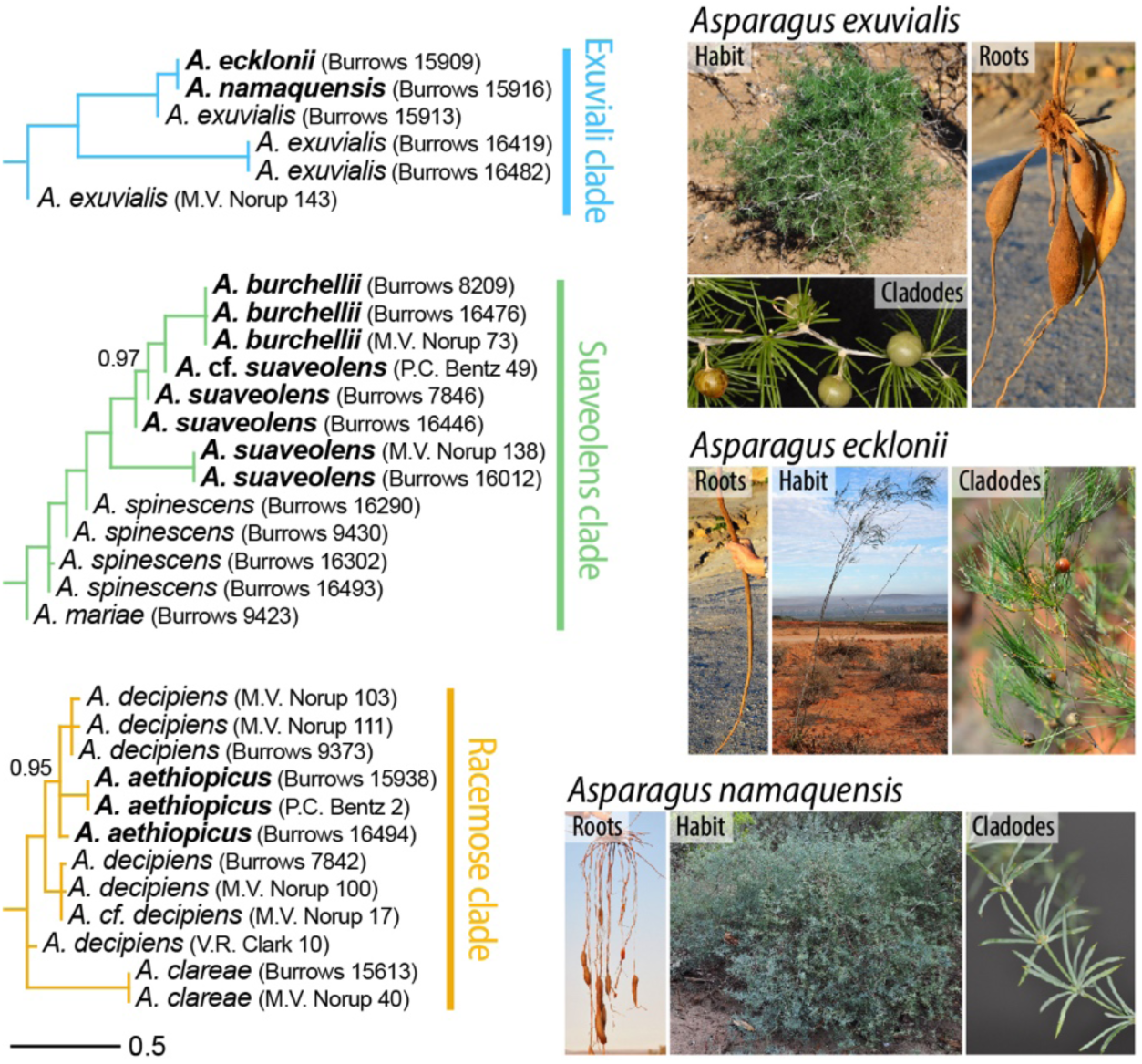
Three subtrees illustrating paraphyly caused by budding speciation from within extant ancestral species of *Asparagus*. Species images show contrasting phenotypes of two budded species (*Asparagus ecklonii* Baker and *Asparagus namaquensis* MS) that evolved from an ancestral population of *Asparagus exuvialis* Burch. in the Exuviali clade. Branch lengths/scale bar correspond to coalescent time units. Support values are local posterior probabilities shown when ≤0.99. Bolded tip labels refer to the putatively budded species. Species images are excerpts from a forthcoming monograph and taxonomic revision of *Asparagus* taxa in southern Africa (J.E. Burrows & S.M. Burrows, in prep.).

Whether driven by geographic isolation or niche specialization (28), budding speciation giving rise to phylogenetically distinct species nested within extant paraphyletic species may be common in plants (29). Three clear-cut examples of possible budding speciation in *Asparagus* were identified in the Racemose, Suaveolens, and Exuviali clades (Fig. 4). Interestingly, there was no strong evidence of interspecific gene flow or any departure from the multispecies coalescent model (71, 72, 73) at any of the nodes (all QD>0.70) shown in Fig. 4, suggesting establishment or initiation of reproductive barriers between ancestral and budded species.

In the Racemose clade, short and largely polytomous branches among paraphyletic accessions of *Asparagus aethiopicus* L. and *Asparagus decipiens* (Baker) MS suggest that *A. aethiopicus* budded from the extant ancestral species *A. decipiens* (Fig. 4). The ranges of these two species are largely disjunct, implicating geography as a reproductive barrier: *A. aethiopicus* is mainly confined to the coastal belt of the Western Cape of South Africa, whereas *A. decipiens* occurs throughout much of the drier regions of the Eastern Cape and eastern reaches of the Western Cape of South Africa (Burrows & Burrows, in prep.). Although hybridization potential between these two species is unknown, the patterns listed above are nonetheless suggestive of an ongoing speciation process.

In the Suaveolens clade, multiple species gradients (i.e., branching events) among accessions of *Asparagus spinescens* Steud. ex Schult. & Schult.f., *Asparagus suaveolens* Burch., and *Asparagus burchellii* Baker (Fig. 4) supports the notion that speciation is not some discrete event that always results in the emergence of bifurcating lineages and extinction of the MRCA, but rather speciation occurs on a continuum and multiple budded species can evolve simultaneously from various ancestral populations (e.g., sympatric formation of multiple host races and species in herbivorous insects) (30). Interestingly, *A. spinescens* is more geographically restricted compared to the two budded species, but it remains possible that the budding of *A. suaveolens* occurred in a small, narrowly distributed population of *A. spinescens*. On the other hand, *A. suaveolens* exhibits one of the widest distributions of African *Asparagus* species and is more widespread than the budded *A. burchellii* (26).

In the Exuviali clade, *Asparagus exuvialis* Burch. —a species that extends widely across South Africa, Namibia, Botswana, southern Zimbabwe, and southern Mozambique (26)—is paraphyletic with two budded species (*Asparagus ecklonii* Baker and *Asparagus namaquensis* MS) that arose from an ancestral *A. exuvialis* (Fig. 4). *Asparagus ecklonii* and *A. namaquensis* are broadly sympatric and narrowly distributed in the Northern Cape and Western Cape of South Africa, each preferring distinct dominant soil types: in the north of its range *A. namaquensis* prefers red Aeolian sands, whereas *A. ecklonii* prefers deep sandy soils in shrublands often dominated by *Restio* (Burrows & Burrows, in prep.). Although the extant ranges of *A. exuvialis*, *A. ecklonii*, and *A. namaquensis* overlap in the Western Cape region, each species exhibits strikingly different growth habits among several other distinct phenotypes (Fig. 4) suggesting realization of different ecological niches in the budded species.

Two extremely polymorphic species in the Myrsiphyllum clade—*Asparagus ovatus* T.M.Salter and *Asparagus asparagoides* (L.) Druce—are both highly polyphyletic in a fashion that is not associated with geography or intraspecific phenotypic variation (Fig. S1). The polyphyly among these Myrsiphyllum lineages may be indicative of a larger taxonomic problem involving the lumping of several cryptic species. Alternatively, *A. asparagoides* and *A. ovatus* may represent “ochlospecies” that formed through a process of repeated isolations and refusions among divergent populations driven by climatic shifts during the Pleistocene (14, 27), or by rapid range expansion and colonization of new habitats without an allopatric phase, leading to a complex of variable intraspecific phenotypes/genotypes (22). Interestingly, two accessions of *A. asparagoides* (*R. Boon 162* & *163*) were sampled from different populations invasive to Australia (Appendix S1) and were placed in disjunct clades within Myrsiphyllum (Fig. S1); suggesting at least two separate introductions to Australia—each likely involving progenitors from the Western Cape of South Africa. Polyphyly was also observed among other taxonomically troublesome species including *Asparagus setaceus* (Kunth) Jessop and *Asparagus densiflorus* (Kunth) Jessop (Results S1), further illustrating the need for continued taxonomic and population-level work to better delimit species within the genus and elucidate the speciation process.

### Major clades of *Asparagus*

Previously, six major clades were hypothesized to encapsulate all *Asparagus* species diversity (12), including the following labeled clades: Racemose, Africani-Capenses, Asparagus, Lignosus, Setaceus, and Myrsiphyllum clades. Those six major clades were largely reinforced by morphological synapomorphies and biogeography (12) and plastome sequences (14). The species tree analysis in this study (Fig. S1) also strongly supported those six previously defined clades (12, 14). However, increased species sampling in our analyses enabled splitting of the former Africani-Capenses clade into four distinct clades and allowed us to test taxonomic hypotheses generated over 100 years ago (Results S1; Table S3). For instance, subgeneric taxon groups proposed by Obermeyer & Immelman (26) were generally more congruent with the 12 major clades in Fig. 1 compared to those hypothesized by Baker (31) and Jessop (48) (Table S3). Our analyses also revealed congruence between the former Capenses and Lignosus clades from Norup et al. (12) and the taxonomic series’ Suaveolens and Retrofracti, respectively, of Obermeyer & Immelman (26); therefore, we refer to those clades as such in this study (Table 1; Fig. 1). The substantial species sampling in our analyses enabled delineation of an additional four major clades in *Asparagus* (i.e., Macowanii, Nutlet, Scandens, and Sympodioidi) which were defined based on phenotypic synapomorphies and strong support for monophyly (Results S1).

We found strong support for a sister relationship between the Asparagus-Exuviali clade and the rest of the genus (Fig. 1), whereas previous studies either weakly (12) or strongly (14) suggested that the Setaceus clade is sister to the rest of *Asparagus* and that the Asparagus-Exuviali clade is more closely related to the Racemose clade (12, 14). Our analysis strongly supports a sister relationship between the Setaceus clade and an Africani-Sympodioidi-Suaveolens-Nutlet clade, which together form a larger clade that is equally related to three other African (bisexual) clades (see backbone polytomy with four leaves in Fig. 1). Bifurcations among the other major clades in *Asparagus* mostly resulted in poor support in previous studies, aside from moderate support for close relations between Suaveolens and Africani (12, 14).

Myrsiphyllum was the only major clade in our analyses with LPP<1.0 which is in-line with previous findings (12) in that inclusion of *Asparagus fasciculatus* Thunb. was weakly supported (Fig. S1). It is possible that *A. fasciculatus* is not in fact part of the Myrsiphyllum clade, but rather a distinct lineage arising from a polytomous node subtending i) the core Myrsiphyllum clade, ii) *A. fasciculatus,* and iii) the Scandens clade; however, that hypothesis was rejected in a polytomy test (Fig. S3). Further, *A. fasciculatus* exhibits diagnostic traits typical of Myrsiphyllum, including connivent filaments and perianth segments (24). The ancestral flower form for the Myrsiphyllum clade likely included those diagnostic floral traits, which are ubiquitous across the clade, except in the species *Asparagus ramosissimus* Baker which exhibits spreading perianth segments and filaments (Results S1). *Asparagus scandens* (Thunb.) Oberm. of the Scandens clade is another taxonomically troublesome species and, due to multiple shared traits (e.g., flattened, leaflike cladodes) *A. scandens* is often lumped into Myrsiphyllum (formerly genus *Myrsiphyllum* Willd.) (24). However, unlike its sister lineage, *Asparagus mollis* (Oberm.) Fellingham & N.L.Mey., *A. scandens* has spreading filaments and perianth segments (24). Due to shared morphology and weak support in phylogenomic analyses (LPP=0.83), it is possible that the Scandens clade is sister to Myrsiphyllum, but our results failed to show statistical support for that hypothesis (Fig. S3). According to our analyses, the most probable relationship between the Myrsiphyllum and Scandens clades was a polytomy including two additional major clades (Fig. 1).

### Origin of cultivated garden asparagus and implications for breeding

Garden asparagus (*A. officinalis*) was cultivated ∼2000–2500 years ago (40) and has since been widely naturalized around the globe (1), resulting in dubious reports of its ‘natural’ range (41) which remains unknown. However, studies have shown that contemporary garden asparagus cultivars significantly differ genetically from putative wild populations in Turkey and Iran (42, 43), suggesting a center of origin in western Asia. To test the geographic origin of *A. officinalis*, we used the extant ranges of its closest relatives to estimate the ancestral, pre-domestication range of the domesticated species. *Asparagus persicus* Baker, *Asparagus pseudoscaber* Grecescu, and *A. officinalis* form a well-supported polytomy (garden asparagus clade: Table 2) sister to *Asparagus palaestinus* Baker (Fig. 2). The estimated ancestral range of the garden asparagus clade was the western Asia/Medit. Basin region (100% probability: Table 2), which is where the closest extant relatives of *A. officinalis* occur and is generally in-line with previous hypotheses that also implicate the Caucasus region as a possible center of origin (44, 45). A greater understanding of interspecies genetic relatedness is crucial to focused breeding efforts aimed to improve the economically important *A. officinalis*, due to its relatively low genetic diversity in breeding programs (69) and absence of valuable agronomic traits that are more common in wild *Asparagus* (e.g., disease resistance and salt, drought, and acid soil tolerance) (70). The robust species tree inference from this study will contribute to future breeding efforts, either for vegetables (e.g., *A. officinalis*) or herbal medicinal purposes (e.g., *Asparagus racemosus* Willd. and *A. cochinchinensis*) (2), by lending insight into divergence of candidate material for interspecific crosses.

## Materials and Methods

### Sample preparation and sequencing

Samples were prepared for Hyb-Seq experiments with Asparagaceae1726 using the exact methods as described by Bentz & Leebens-Mack (52). In sum, DNA was extracted from silica dried, flash frozen, or herbarium voucher tissue and Illumina DNA-sequencing libraries were prepared with universal Y-yoke stub adapters and dual-indexed iTru primers (46) aiming for an average fragment length of 350–550 bp. Genomic libraries were pooled for hybridization with Asparagaceae1726 probes (52), then captured target DNA fragments were enriched for 14 cycles of PCR (52). Additional sample and data generation details can be found in Appendix S1.

### Species sampling

We sampled as many species possible of *Asparagus*, attempting to include species spanning the entire geographic range of the genus, as well as one *Hemiphylacus*, and three outgroups from different Asparagaceae subfamilies (Table 1). Accessions for several undescribed taxa were included in our dataset (Table 1), to be described in a forthcoming monograph and revision of the genus (Burrows & Burrows, in prep.), and are labeled with ‘MS’ (Appendix S1). All major clades of *Asparagus* were sampled in this study, including several DNA accessions from Norup et al. (12) (Appendix S1). For DNA isolation, tissue was collected from wild populations, herbarium vouchers, or cultivated plants growing in greenhouses or botanical gardens (Appendix S1). We aimed to sample multiple individuals from different populations of each species when possible. Ten accessions were from whole genome shotgun sequencing (WGS) experiments repurposed for this study (Appendix S1).

### Target ortholog and paralog assembly

Lingering adapter sequences were removed from reads, mismatched base pairs were corrected, and reads shorter than 21 bases were filtered out using fastp v.0.23.2 (49). The HybPiper v.2.1.6 pipeline (23) was then used to i) map the filtered reads to target nucleotide sequences with BWA-MEM (50), ii) assemble mapped reads into contiguous sequences with SPAdes (51), iii) extend gene assemblies into flanking intronic regions, and iv) test for paralogs with default HybPiper parameters. BLASTX was used for read alignments with HybPiper for some WGS accessions (Appendix S1). The Asparagaceae1726 v.1.1 (https://github.com/bentzpc/Asparagaceae1726) target sequence file was used as reference for HybPiper processing, as performed by Bentz & Leebens-Mack (52).

### Phylogenomic analysis

Multiple sequence alignments (MSAs) for each target locus were produced using MAFFT v.7.487 with the flag *–auto* (53). Poorly aligned sequences were trimmed using trimAl v.1.4.1 with the flag *-automated1* (54). IQ-TREE v.1.6.12 (55) was used for Maximum Likelihood (ML) gene tree inference with 1000 ultrafast bootstraps (BS) and the *MFP* option which allows ModelFinder (56) to choose the best fit substitution model based on Bayesian information criterion. Species tree inference was derived from the full collection of unrooted ML gene trees using wASTRAL-unweighted v1.16.3.4 (57) by optimizing the objective function of ASTRAL-III v.5.7.8 (58), which employs a coalescent-based summary method that is statistically consistent under the multi-species coalescent model. Prior to the ASTRAL analysis, gene tree branches with <10% BS support were collapsed into polytomies to improve the accuracy of species tree inference (58). The resulting ASTRAL tree was rooted with outgroup taxa (Table 1) then a polytomy test was performed with ASTRAL-III; species tree nodes were subsequently collapsed when a polytomy could not be rejected (*p*-value >0.05). Along with metrics reported by ASTRAL, gene tree quartet support (e.g., QD) was also assessed with Quartet Sampling v.1.3.1 (47), and RAxML-ng v.1.2.2 (25) as the likelihood evaluation engine, across all nodes. Quartet Sampling was performed with 1000 replicates and a likelihood threshold of 2.0 (47). Coalescent-based, rather than concatenation, methods were used to analyze nuclear genes because concatenation implicitly assumes an absence of recombination and ILS (i.e., a single history is shared amongst all genes) and can result in statistical inconsistencies in multi-locus datasets with high levels of gene tree discordance (59, 60); whereas coalescent approaches assume free recombination between genes, accounting for gene tree–species tree discordance due to ILS.

### Ancestral range estimation

A second species tree reconstruction was performed with less terminal branches for ancestral range estimations by forcing closely related accessions together in quartet analyses of a subsequent ASTRAL-III analysis, which allows pre-defined lineages to coalesce into a single, terminal branch (58). Single branch coalescence enables terminal branch length estimation, representing coalescent time units. Accessions were forced into a single lineage/branch only if they formed a well-supported clade (LPP ≥0.98 in the previously inferred ASTRAL tree) and exhibited overlapping geographic distributions (Appendix S1). For instance, lineages with a narrow geographic distribution nested in an otherwise broadly distributed clade/range were collapsed into a single branch. Based on existing literature of *Asparagus* biogeography (1, 12, 14) and the biologically informative geographic areas defined by Taxonomic Databases Working Group (68), we defined 11 areas (states) for ancestral range estimation using stochastic mapping with BioGeoBEARS v.1.1.1 (61, 62). We analyzed and chose the best of three biogeographic models implemented with BioGeoBEARS: i) the DEC model (63), ii) an ML implementation of the Dispersal-Vicariance Analysis (64) (termed DIVALIKE), and iii) an ML implementation of BayArea (65) (termed BAYAREALIKE) (62, 66). We also tested whether allowing for founder-events within clades (parameter *j*) increased model fit, based on likelihood-ratio tests for each nested model, considering a *p*-value <0.05 as significant. The best fit model was chosen based on Akaike Information Criterion (AIC), for which we calculated sample size corrected AIC (AICc) then AIC weights (AICc_wt) to represent the relative likelihood of each model, choosing the model with the highest (best) AICc_wt score (67). Each biogeographical test was unconstrained, allowing for equal probabilities among all possible area combinations for dispersal routes. Defined geographic areas included Southwestern + Central Europe; Eastern Europe + Caucasus; Northern Africa + Macaronesia; Northern Tropical Africa; Central Tropical Africa; Central Asia; Southern Africa + Madagascar; Western Asia to Medit. Basin; Arabian Peninsula; Eastern Asia to Malesia; and the Indian Subcontinent (Fig. S8). Prior to analysis with BioGeoBEARS, a subset of noninformative branches were pruned from the input ASTRAL tree to make the analysis computationally tractable (Appendix S1).

## Data availability statement

All relevant result files and original scripts from this study are available at https://zenodo.org/doi/10.5281/zenodo.10804898. Sequencing reads from this study were deposited to NCBI Sequence Read Archive under the BioProjects PRJNA1088837 and PRJNA1088858.

## Acknowledgments

This work was supported by the US National Science Foundation (DEB-2110875). We thank the following kind people for aiding in sample collection: Miguel Ángel González Pérez (Jardín Botánico Canario Viera y Clavijo, Gran Canaria, Spain); Zach Stansell (USDA-ARS, Geneva, New York, USA); Stéphane M. Bailleul, Frédéric Coursol, and team (Espace pour la vie, Jardin botanique de Montréal, Canada); Mark Weathington (JC Raulston Arboretum, Raleigh, North Carolina, USA); Tony Avent, Zac Hill, Patrick McMillan, and Amanda Wilkins (Juniper Level Botanical Garden, Raleigh, North Carolina, USA); Mason McNair (Clemson University, Florence, South Carolina, USA); Carlos Lobo (Jardim Botânico da Madeira, Instituto das Florestas e Conservação da Natureza, Madeira, Portugal); Sean C. Lahmeyer (The Huntington Botanical Gardens, San Marino, California, USA); Peter Brownless (Royal Botanical Gardens, Edinburgh, Scotland); Craig Jackson (The John Fairey Garden Conservation Foundation, Hempstead, Texas, USA); Maria Norup; Laura Genco and Davide Bonaviri (WWF Italy, Capo Rama reserve); Francesco Mercati (Università Mediterranea degli Studi di Reggio Calabria, Italy); and Akira Kanno (Graduate School of Life Sciences, Tohoku University, Japan).

## Supporting Information

**Appendix S1:** Accession and species sampling information

**Appendix S2:** Target capture results reported by HybPiper

**Appendix S3:** Ancestral range probabilities for all branches/nodes reported by BioGeoBEARS

Appendices S1–S3 are available at https://zenodo.org/doi/10.5281/zenodo.12812248.

### This PDF file includes

Results S1

Figures S1 to S8

Tables S1 to S3

SI References

## Results S1

### Sequencing and target capture

After minimal filtering, the total number of Hyb-Seq reads per sample were variable, ranging from 162,802 to 174,960,651 (mean=33,736,507; median=27,672,928). The fraction of reads that mapped to reference target sequences (percent on-target reads) were also variable among Hyb-Seq experiments, ranging from 3.1% to 77.4% (mean=39%; median=40%). All samples (Hyb-Seq and whole genome shotgun sequencing [WGS]) yielded >1300 target orthologs, aside from three: *A. secalemontanus* (*Burrows & Burrows 15560*) yielded 483, *A. denudatus* (*Burrows & Burrows 8392*) yielded 711, and *A. angusticladus* (*Burrows & Burrows 12678*) yielded 973 (Appendix S2). Only 14 Hyb-Seq accessions yielded all 1726 targets, however 165 accessions yielded 1720–1726 targets, 210 yielded 1700–1726 targets, and 258 yielded 1650–1726 targets (Appendix S2). Eight of the ten WGS samples yielded >1700 targets (Appendix S2).

### Putative paralogs captured by Asparagaceae1726

Potential paralogs were scored for each locus using default HybPiper (13) run parameters, which report paralogs based on the coverage and number of SPAdes (20) contig assemblies mapping to a target sequence with inferred paralog scoring based on “depth” (> 75% of target length covered by > 2 shorter contigs in addition to the putative ortholog assembly) and contiguous “length” (> 1 SPAdes contig mapping to > 75% of the reference target sequence) of non-primary assemblies for each target (13). Target paralogs detected by HybPiper’s “length” criteria were generally low in this dataset, ranging between 5–33 (mean/median=15) in WGS samples and 0–231 (mean=16; median=15) in Hyb-Seq samples (Appendix S2). The *Hemiphylacus* sample yielded the largest number of possible long paralog assemblies—231 in total—whereas the next two highest totals were 97 (in *A. gobicus* [P.C. Bentz 75]) and 50 (in *A. niassanus*) (Appendix S2). Compared to the long paralog warnings, HybPiper flagged an average of approximately three times as many paralogs with its “depth” detection scheme per sample (Appendix S2).

### Rampant paraphyly and polyphyly across the *Asparagus* phylogeny

Across the *Asparagus* tree, there are many examples of polyphyletic species including strongly supported branches (local posterior probability [LPP]=1). In the Racemose clade, *A. decipiens* was paraphyletic with the budded species *A. aethiopicus* (Fig. S1); *A. tugelicus* evolved in a paraphyletic clade of *A. kwazuluanus*; *A. filicladus* evolved from an ancestral form of *A. acocksii*, which was paraphyletic with the former; and *A. densiflorus* was polyphyletic with two cultivated lineages forming a clade separate from wild forms of the species (Fig. S1). *Asparagus densiflorus* (Kunth) Jessop has long been plagued by taxonomic issues (1). Both *A. densiflorus* clades may represent distinct species, or perhaps the *A. densiflorus* cultivars sampled here were derived from separate ancestral species not sampled in this study. Resolving either of those hypotheses is out of the scope of this study, but the polyphyly of *A. densiflorus* in the current analysis illustrates a need for continued taxonomic work on species circumscription within the genus. For instance, recent taxonomic work (Burrows & Burrows, in prep.) shows that *A. densiflorus* of Obermeyer & Immelman (2) can be divided into nine clearly separable species. In the Setaceus clade, *A. setaceus* is polyphyletic, arising once in a clade with *A. sylvicola* and again in a clade with *A. brevipedicellatus* (Fig. S1). *Asparagus setaceus* (Kunth) Jessop is another taxonomically problematic species and was polyphyletic in our species tree analysis (Fig. S1). However, one of the two clades of *A. setaceus*, including an accession from Li et al. (3) and *Burrows & Burrows 16255*, may represent *Asparagus plumosus* Baker—a species now synonymous with *A. setaceus* (4). In the Myrsiphyllum clade, accessions of *A. asparagoides* and *A. ovatus* were highly polyphyletic and paraphyletic with various species (Fig. S1). In the Suaveolens clade, *A. spinescens* was paraphyletic with the budded species *A. suaveolens* and *A. burchelii*; *A. flavicaulis* was paraphyletic with *A. candelus*; and *A. capensis* was either paraphyletic with *A. stipulaceus*, *A. litoralis*, and *A. praetermissus,* or polyphyletic (Fig. S1). In the Africani clade, *A. lugardii* was polyphyletic due to one accession (i.e., *Burrows & Burrows 15214*) residing outside a clade with the remaining seven accessions of the species (Fig. S1). In the Exuvialis clade, *A. exuvialis* was paraphyletic with two budded species: *A. ecklonii* and *A. namaquensis* (Fig. S1). In the Asparagus clade, a grade of *Asparagus gobicus* N.A.Ivanova ex Grubov accessions caused this species to be paraphyletic with the nested/budded species *Asparagus dauricus* Fisch. ex Link and *Asparagus kiusianus* Makino. All three of these species occur in similar environments – arid, sandy wastelands sometimes by the sea – but exhibit different geographic ranges: *A. kiusianus* is endemic to the Japanese islands (18); *A. dauricus* is widespread across Mongolia, China, and Korea, and the extant ancestral species *A. gobicus* is more restricted to Mongolia and parts of China (19). Also in the Asparagus clade, *A. longiflorus* resulted in a polytomy with *A. breslerianus*; and *A. persicus* and *A. pseudoscaber* were paraphyletic and in a polytomy with *A. officinalis* (Fig. S1).

### Range expansions out of Africa

We found support for eight independent shifts out of Africa that led to colonization of Eurasia, six of which involved the most recent common ancestor (MRCA) of extant bisexual lineages, including: the MRCA of *A. cooperi* and *A. devenishii* (Africani clade); *A. africanus* (Africani clade); *A. virgatus* (setaceus clade); *A. falcatus* (Racemose clade); *A. racemosus* (Racemose clade); and *A. abyssinicus* (Asparagus clade). We also found support for three separate expansions into Macaronesia involving two bisexual lineages that dispersed from southern Africa (i.e., the MRCA of *A. umbellatus* and *A. arborescence* [Setaceus clade] and the MRCA of *A. scoparius* and *A. plocamoides* [Africani clade]) and the dioecious species *A. horridus* (Asparagus clade) which likely dispersed from the Arabian Peninsula (Fig. S6a) or northern Africa (Figs. S6b, S7). Following the establishment of the Eurasian dioecy clade in eastern Asia, a secondary expansion led to colonization of Europe (main text, Fig. 3), which disagrees with previous findings that suggest dispersal into Eurasia began westward from southern Africa to northern Africa and Europe, then eastward into Asia (12). However, the sparse sampling of dioecious species by Norup et al. (12), and application of misreported range data for *A. officinalis* (further discussed in main text), likely explains our incongruent results.

### Delineation of four additional major clades of *Asparagus*

#### Scandens clade

*Asparagus scandens* (Thunb.) Oberm. of the Scandens clade has long troubled taxonomists and, due to the sharing of several traits typical of Myrsiphyllum species—including flattened (leaflike) cladodes—*A. scandens* is often lumped into Myrsiphyllum (formerly genus *Myrsiphyllum* Willd.) (14). The key traits commonly used to identify Myrsiphyllum species are the presence of basally connivent perianth segments and connivent filaments (15), but this would exclude the Scandens clade (as delimited here) as well as *Asparagus ramosissimus* Baker (14) for which there was strong support for its placement in the Myrsiphyllum clade (Fig. S1). Not only does the Scandens clade exhibit morphological affinity to the Myrsiphyllum clade, but phylogenomic analysis supports a Myrsiphyllum-Scandens clade (LPP=0.83) over alternative resolutions (Fig. S5). Due to shared morphology and weak support in phylogenomic analyses it is possible that the Scandens clade is more related to Myrsiphyllum than any other clade, but our analyses failed to show consistent support for that hypothesis. According to our analyses, the most probable relationship between the Myrsiphyllum and Scandens clades was a polytomy including two additional major clades (Fig. S1). Considering the absence of a connivent perianth and filaments in *A. ramosissimus*, it is possible that ancestral polymorphisms—both phenotypic and genotypic—were lingering in ancestral populations of *Asparagus* and were carried over into the two contemporary clades Myrsiphyllum and Scandens. For example, perhaps over time, as new species formed and continued to diverge from ancestral populations, various polymorphisms (i.e., connivent or spreading perianth segments and filaments) may have become fixed in different populations/species (i.e., *Asparagus mollis* (Oberm.) Fellingham & N.L.Mey., in the Scandens clade and *A. ramosissimus* in the Myrsiphyllum clade) leading to different ancestral traits sorting between closely related species. Of course, it remains possible that *A. mollis* and *A. ramosissimus* independently evolved floral traits that differ from their ancestors. In any event, the placement of *A. scandens* in our analyses (Fig. S1) is incongruent with previous work that placed it sister to the Retrofracti clade with significantly less sequence data (12).

#### Nutlet clade

Species tree analysis also revealed strong support for a monophyletic Nutlet clade (Fig. S1), suggesting a single origin of a nutlet fruit type in *Asparagus*. A nutlet fruit represents the main synapomorphy of the Nutlet clade and was likely derived from an ancestral black or red berry fruit type, which are ubiquitous in the Suaveolens and Africani-Sympodioidi clades, respectively. Norup et al. (12) sampled two nutlet bearing species and showed moderate support for a bifurcation between those and the Suaveolens clade but referred to the nutlet bearing clade as ‘Recurvispinus’ and lumped it in with the greater Suaveolens clade.

#### Sympodioidi clade

As described here, the Sympodioidi clade is equivalent to the Sympodioidi series previously described (2) and is identifiable by their dichotomously branching stems with hard/woody scales and umbellate inflorescences. The Sympodioidi clade was strongly supported as monophyletic and sister to the greater Africani clade (Fig. S1), whereas a previous analysis of the same Sympodioidi species showed weak support for the nesting of Sympodioidi within the Africani clade (12).

#### Macowanii clade

Resolution of a Macowanii clade sister to the Racemose clade was strongly supported in analyses performed here (Fig. S1) and in previous work (12). The early branching of Macowanii taxa from the larger Racemose clade, coupled with distinct inflorescence morphologies found in Macowanii taxa (i.e., solitary flowers) compared to most Racemose taxa (i.e., compound inflorescences) supports delineation of the two clades.

#### New combinations of *Asparagus* species associated with this study

Alongside this study, a taxonomic revision of *Asparagus* species in Africa was completed by John E. Burrows and Sandra M. Burrows for a separate publication. (Appendix S1).The following species and subspecies (marked with ‘MS’) were described for the first time in the aforementioned revision of *Asparagus*: *Asparagus atrobracteatus* MS, *Asparagus brevipedicellatus* MS subsp. *barbertonicus* MS, *Asparagus brevipedicellatus* MS subsp. *brevipedicellatus* MS, *Asparagus candelus* MS, *Asparagus cederbergensis* MS, *Asparagus cupressoides* MS, *Asparagus dolomiticus* MS, *Asparagus ferox* MS, *Asparagus helmei* MS, *Asparagus humifusus* MS, *Asparagus inopinatus* MS, *Asparagus kwazuluanus* MS, *Asparagus linearis* MS, *Asparagus macrocarpus* MS, *Asparagus namaquensis* MS, *Asparagus niassanus* MS, *Asparagus oubergensis* MS, *Asparagus paniculatus* MS, *Asparagus petiolatus* MS, *Asparagus pongolanus* MS, *Asparagus praetermissus* MS, *Asparagus procerus* MS, *Asparagus pseudoconfertus* MS, *Asparagus recurvatus* MS, *Asparagus saxicola* MS, *Asparagus secalemontanus* MS, *Asparagus soutpansbergensis* MS, *Asparagus tugelicus* MS, *Asparagus zanjianus* MS, *Asparagus lignosus* Burm.f. subsp. *rooibergensis* MS, *Asparagus capensis* L. subsp. *saximaritimus* MS, *Asparagus lignosus* Burm.f. subsp. *rooibergensis* MS.

Additionally, three taxa were raised to the species level and nine were re-instated, since their taxonomic reorganization in Flora of Southern Africa (2), Flora of Tropical East Africa (16), Flora Zambesiaca (17), and Plant of the World Online (4): *Asparagus litoralis* (Suess. & Karling) MS was raised to species from *Asparagus capensis* L. var. *litoralis* Suess. & Karling; *Asparagus decipiens* (Baker) MS was raised to species from *Asparagus racemosus* Willd. var. *decipiens* Baker; *Asparagus comatus* (Baker) MS was raised to species from *Asparagus sarmentosus* L. var. *comatus* Baker. The following taxa were reinstated, relative to previous taxonomic combinations: *Asparagus multiflorus* Baker, *Asparagus puberulus* Baker, *Asparagus abyssinicus* Hochst. ex A.Rich, *Asparagus lugardii* Baker, *Asparagus myriocladus* Baker, *Asparagus sprengeri* Regel, *Asparagus ecklonii* Baker, *Asparagus wildemanii* Weim., *Asparagus zanzibaricus* Baker, *Asparagus racemosus* Willd. var. *zeylanicus* Baker. See Appendix S1 for more information about specific samples and taxonomic authorities used in this study.

---

**Figure S1.**
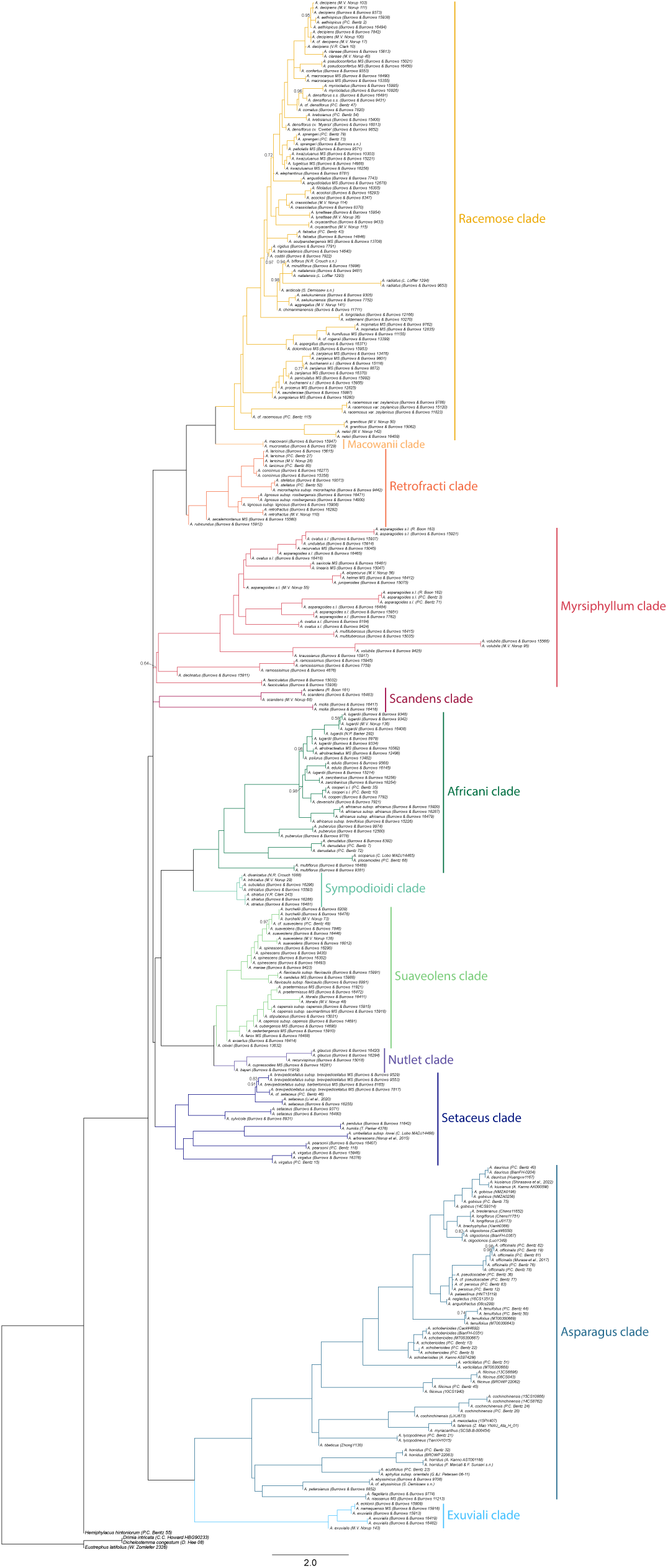
Final species tree phylogram of all accessions from this study showing local posterior probability branch support when <0.99. Bifurcations for which a polytomy could not be rejected (*p*-value >0.05) when tested with ASTRAL (5) were collapsed into polytomies.

**Figure S2.**
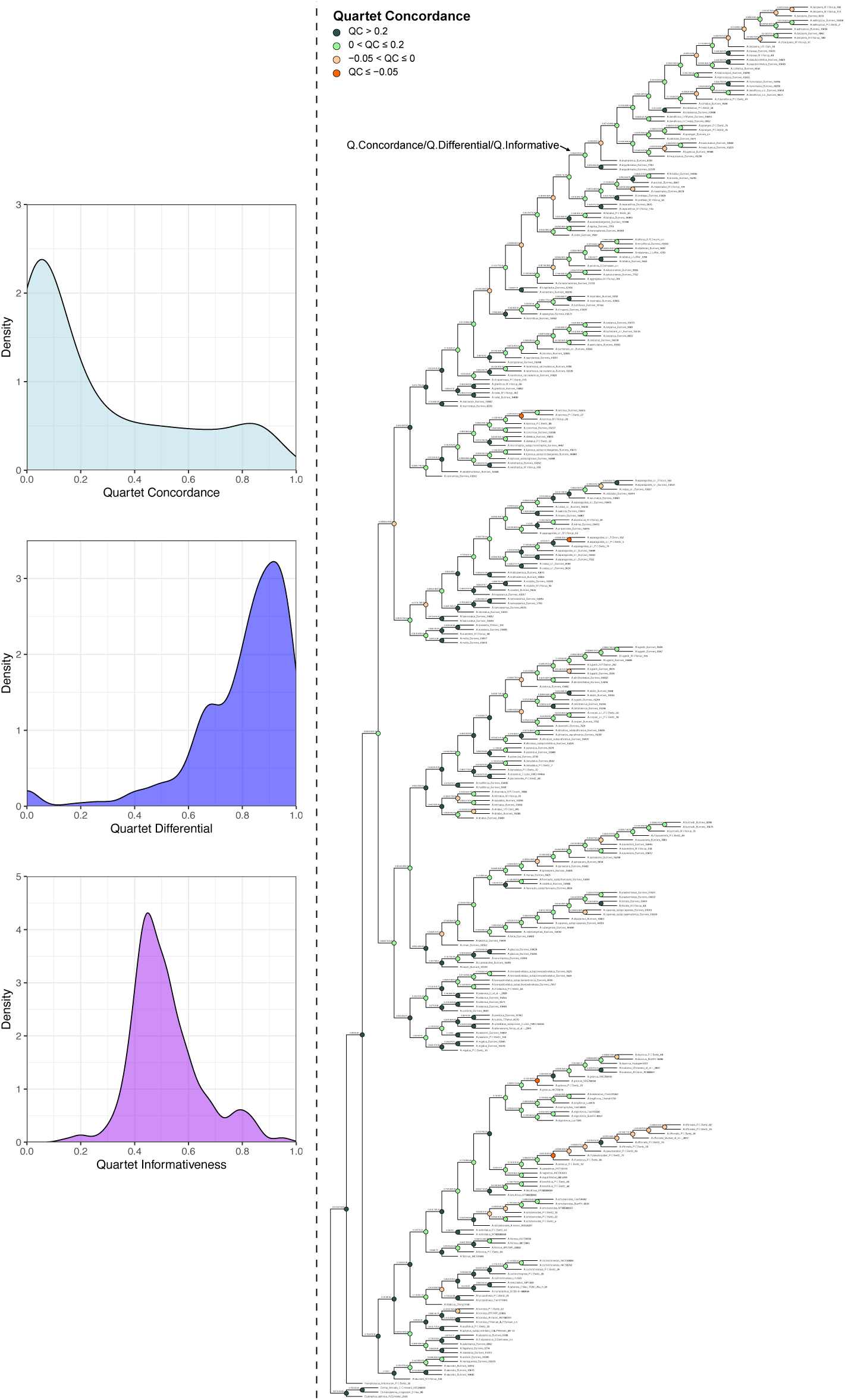
Right panel shows bifurcating species tree cladogram with results from Quartet Sampling (6) (plot: https://github.com/ShuiyinLIU/QS_visualization). Left panel shows histograms of the following metrics from Quartet Sampling. Quartet Concordance overall had a left skewed density distribution (mean=0.23; median=0.09), which is suggestive of considerable gene tree discordance at many nodes. Quartet Differential overall (mean=0.87; median=0.91) suggested limited skew in alternative quartet support. Quartet Informativeness (QI) was moderate across all nodes (mean=0.52; median=0.48) indicating a high degree of information in the gene trees.

**Figure S3.**
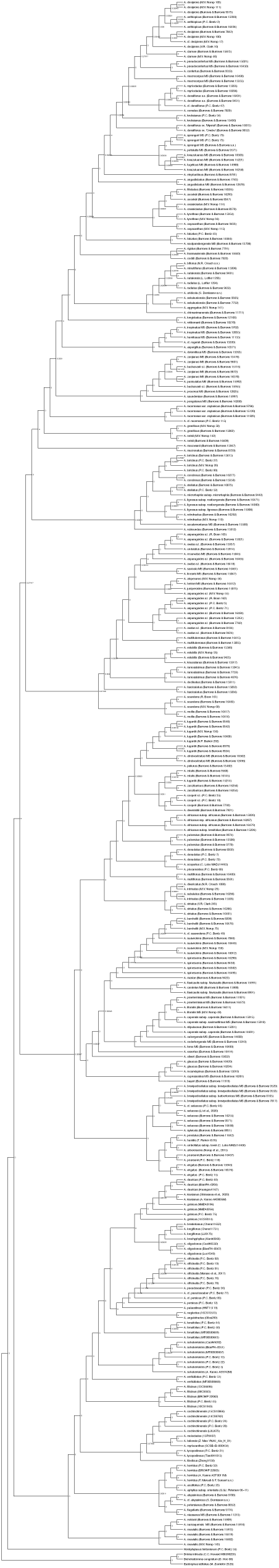
Preliminary species tree cladogram of all accessions showing *p*-values from a polytomy test performed with ASTRAL (5). A polytomy could not be rejected for branches with a *p*-value >0.05 and were subsequently collapsed.

**Figure S4.**
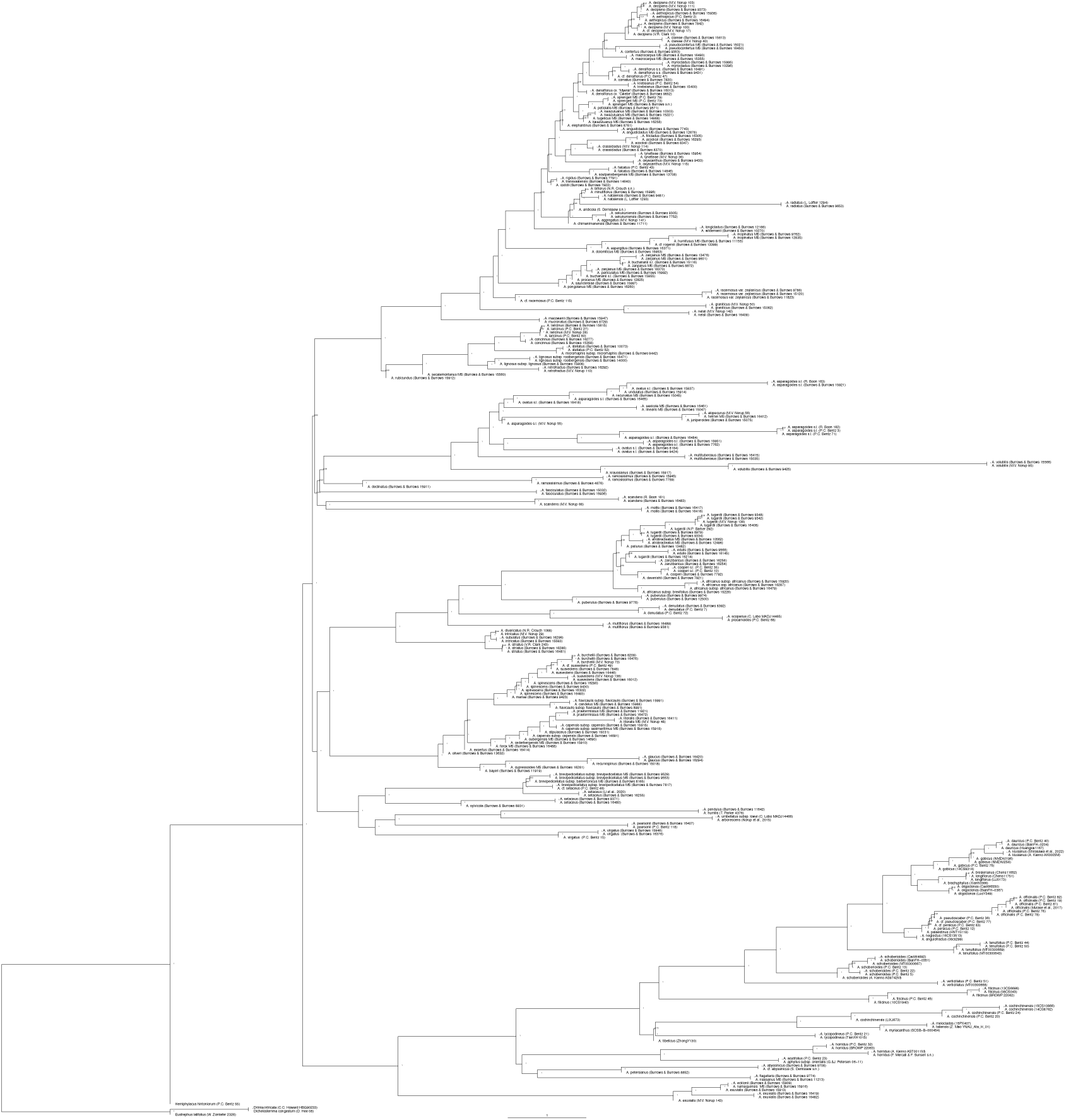
Preliminary species tree phylogram of all accessions showing local posterior probability branch support from ASTRAL (5) for forced bifurcations.

**Figure S5.**
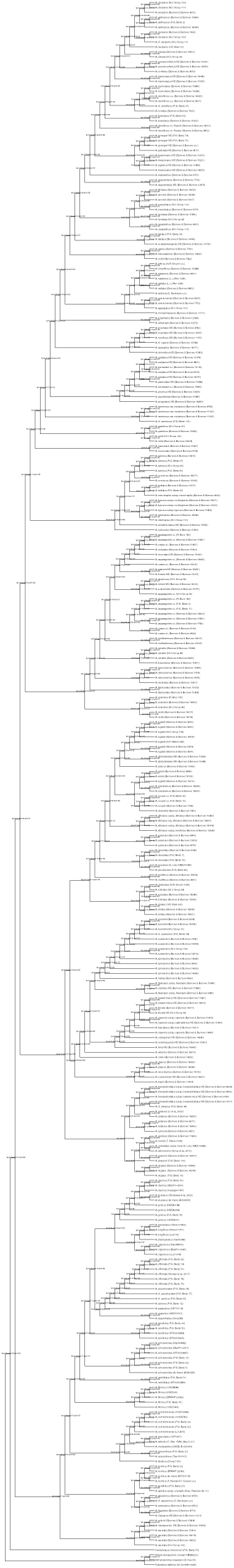
Preliminary species tree cladogram showing quartet frequency support from ASTRAL (5) for the top three topologies. Frequencies represent proportion of gene tree quartets supporting the main topology (*q1*: shown) and two alternatives (*q2* and *q3*).

**Figure S6a.**
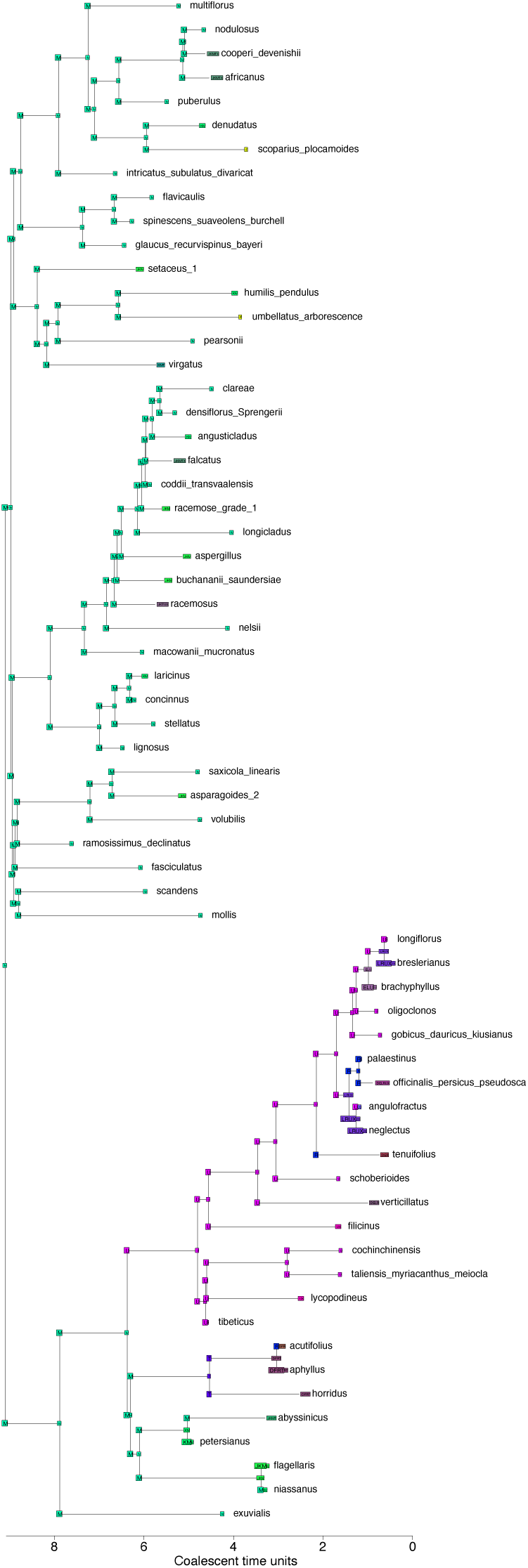
Ancestral range estimation results showing letter codes representing the most probable ancestral *range* for each internal node and split according to BioGeoBEARS model DEC+J (7). Splits are defined as the *ranges* immediately following cladogenesis and are plotted on branch corners. Geographic *areas* include: (D) Southwestern + Central Europe, (E) Eastern Europe + Caucasus, (F) Northern Africa + Macaronesia, (J) Northern Tropical Africa, (K) Central Tropical Africa, (L) Central Asia, (M) Southern Africa + Madagascar, (R) Western Asia to Mediterranean Basin, (T) Arabian Peninsula, (U) Eastern Asia to Malesia, and the (X) Indian Subcontinent. *Area* = discrete geographic region. *Range* = species distribution encompassing any combination of *areas*. See Dataset 3 for a list of collapsed lineages.

**Figure S6b.**
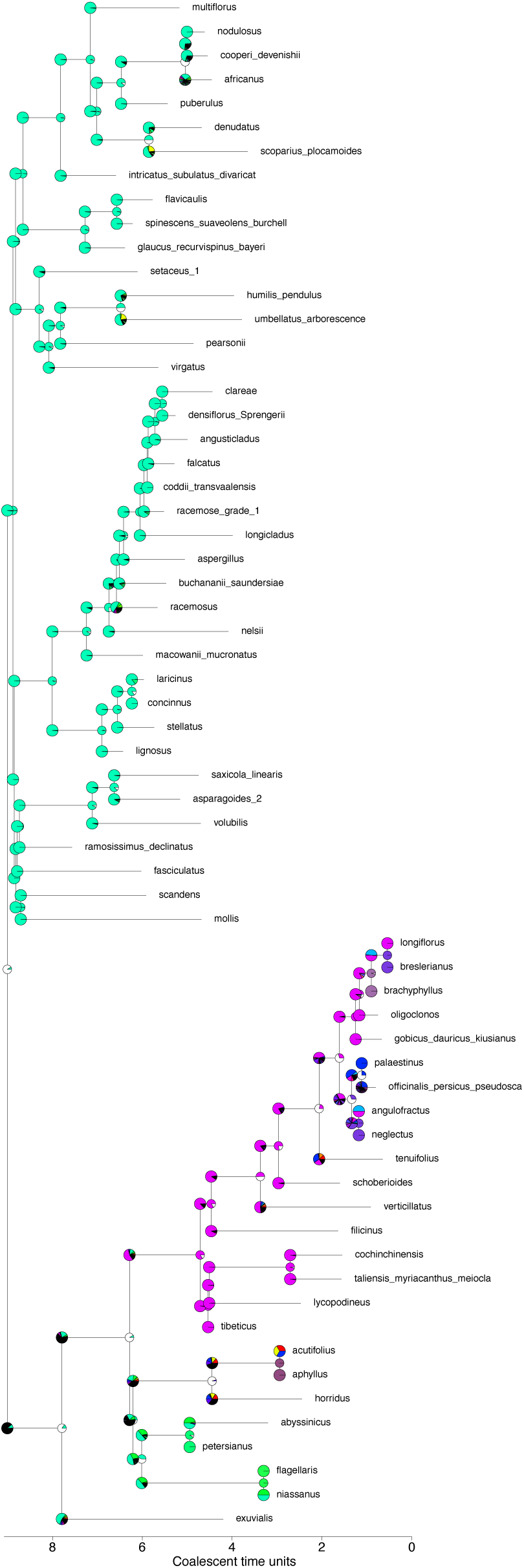
Ancestral range estimation probabilities (pies) for the ancestral *range* distribution of each internal node and split estimated using the BioGeoBEARS model DEC+J (7). Pie colors correspond to geographic *ranges* plotted on the tree in Fig. S6a. Splits are defined as the *ranges* immediately following cladogenesis and are plotted on branch corners. *Area* = discrete geographic region. *Range* = species distribution encompassing any combination of *areas*.

**Figure S7a.**
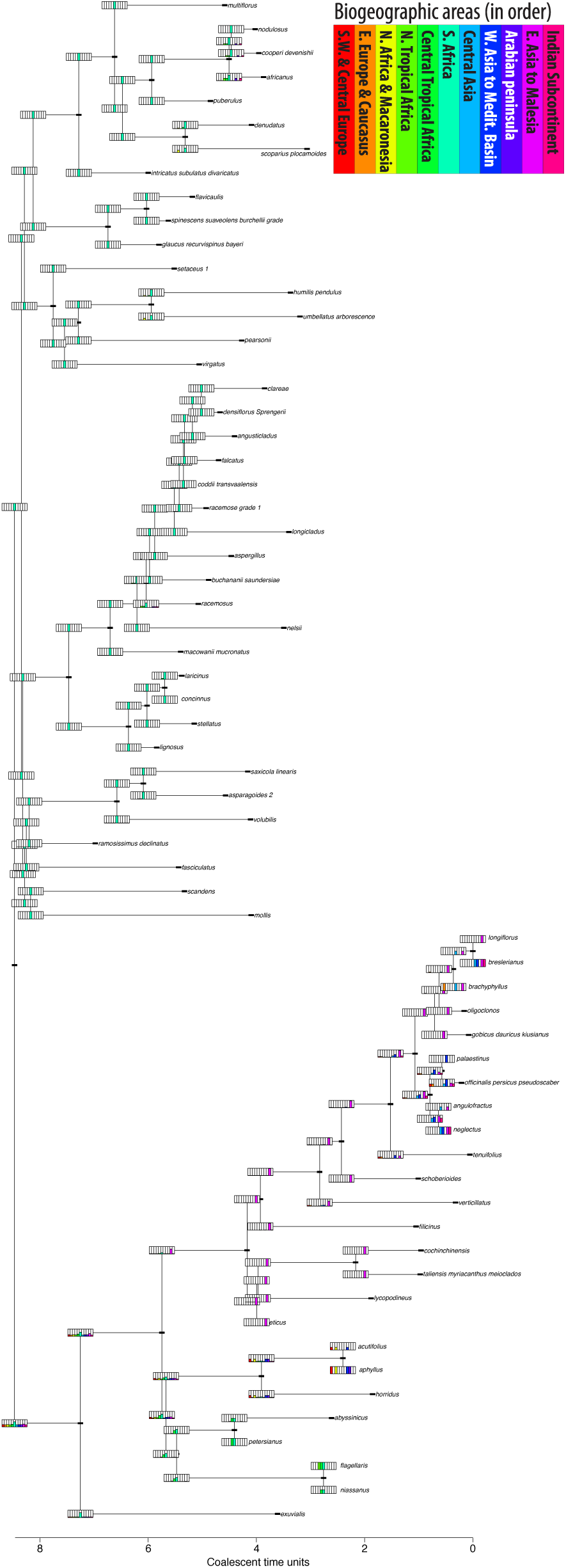
BioGeoBEARS (7) plot showing bar charts of relative probabilities of geographical *area* occupancy (i.e., probability that a *range* included any of the 11 predefined *areas*) for each internal split. Splits are defined as the *ranges* immediately following cladogenesis and are plotted on branch corners. *Area* = discrete geographic region. *Range* = species distribution encompassing any combination of areas.

**Figure S7b.**
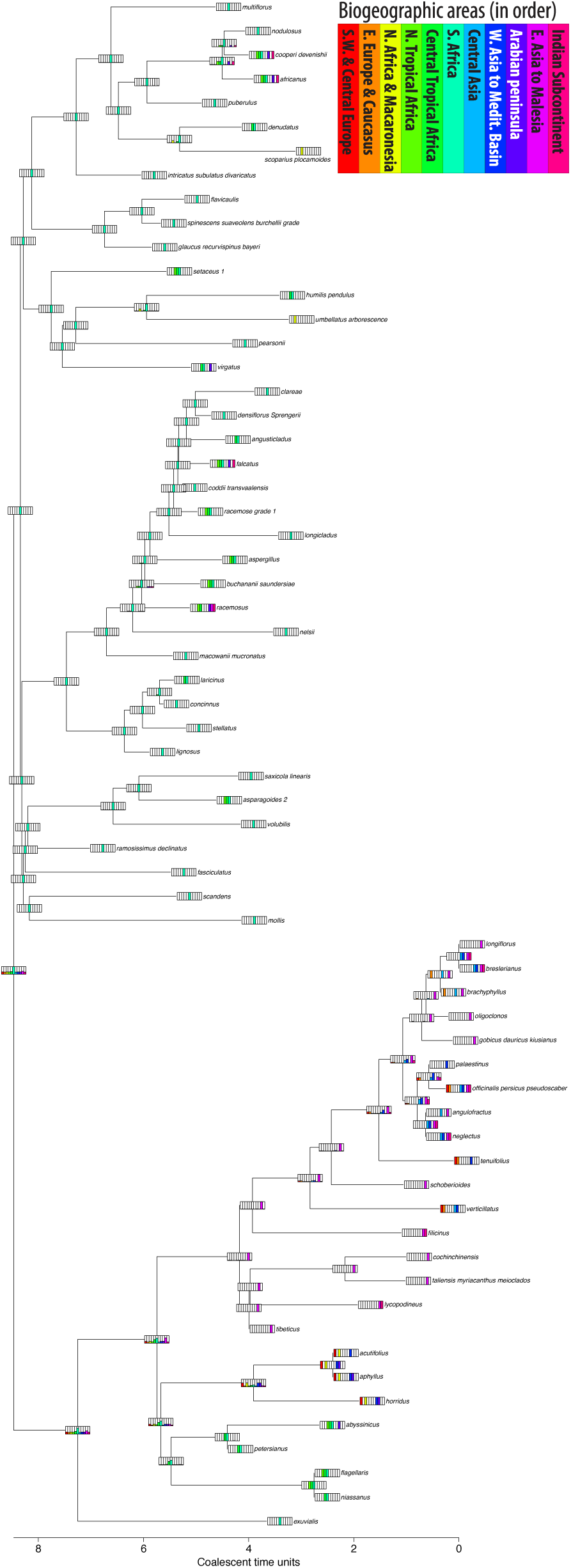
BioGeoBEARS (7) plot showing bar charts of relative probabilities of geographical *area* occupancy (i.e., probability that a *range* included any of the 11 predefined *areas*) for each node. *Area* = discrete geographic region. *Range* = species distribution encompassing any combination of areas.

**Figure S8.**
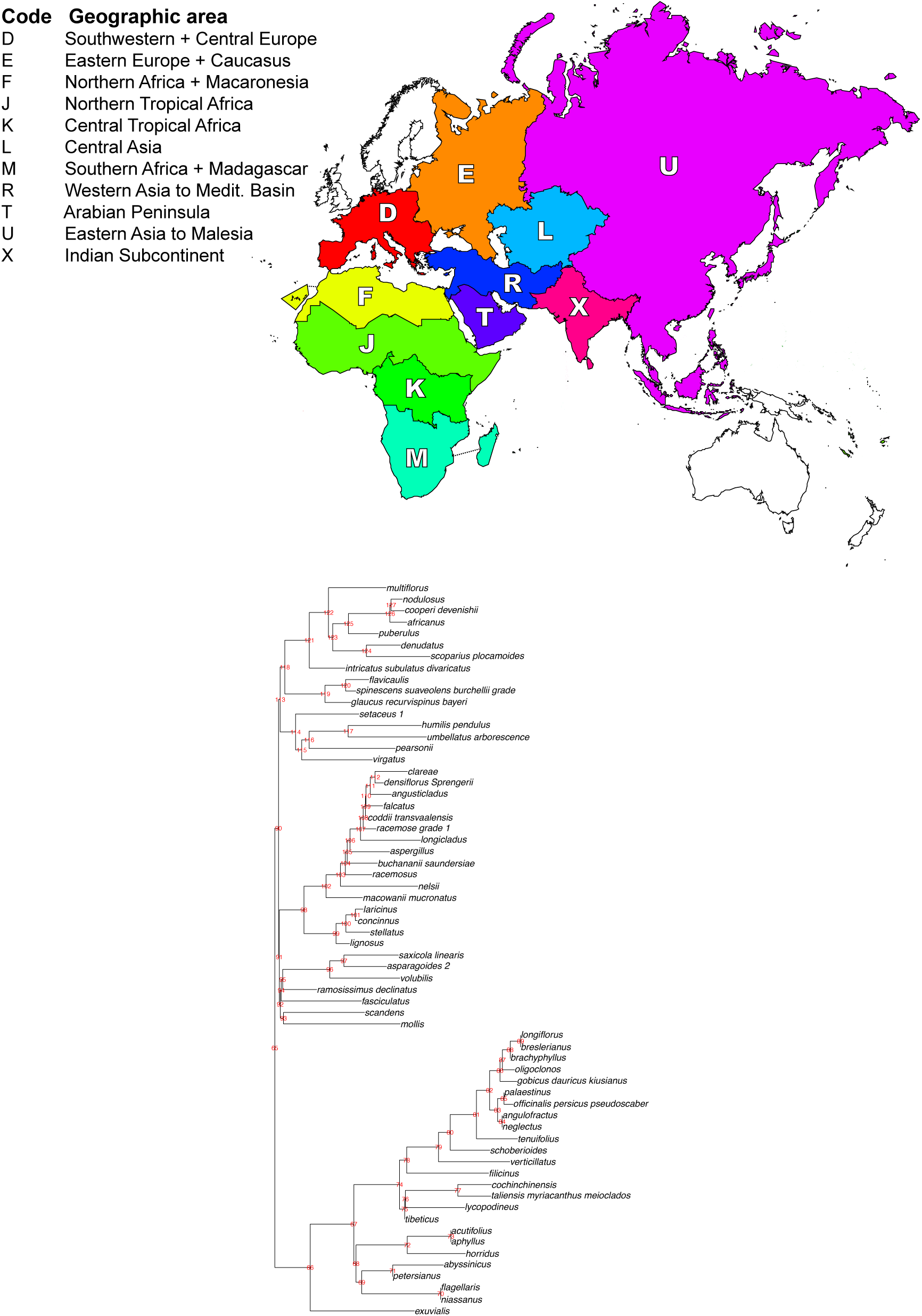
Top plot shows the 11 geographic *areas* defined for ancestral range estimation with BioGeoBEARS (7). Bottom plot shows the phylogeny used for ancestral range estimation with node numbers corresponding to BioGeoBEARS results in Dataset S2. Australia was not included in ancestral range tests since *Asparagus racemosus* Willd. is the only species native there.

**Table S1.** Model test results among biogeographic models estimated with BioGeoBEARS. (**7**).

| Model | LnL | Parameters | Parameter estimates |  |  | AICc | AICc_wt |
| --- | --- | --- | --- | --- | --- | --- | --- |
|  |  |  | *d* | *e* | *j* |  |  |
| DEC | -305.6 | 2 | 0.053 | 0.085 | 0 | 615.4 | 0.2 |
| DEC+J | **-303.1** | **3** | **0.069** | **0.36** | **1.00E-05** | **612.7** | **0.8** |
| DIVALIKE | -361.5 | 2 | 0.06 | 1.00E-12 | 0 | 727.3 | 1.00E-25 |
| DIVALIKE+J | -361.5 | 3 | 0.06 | 1.20E-08 | 1.00E-05 | 729.5 | 3.40E-26 |
| BAYAREALIKE | -321.8 | 2 | 0.04 | 0.2 | 0 | 647.7 | 1.90E-08 |
| BAYAREALIKE+J | -308.6 | 3 | 0.038 | 0.16 | 0.0048 | 623.7 | 0.0033 |

**Note:** AICc = Akaike information criterion (AIC) corrected for sample size. AICc\_wt = AIC weighted based on relative likelihood of the model divided by the sum of the relative likelihood of all models (higher AICc\_wt values are favored) (8). Parameter definitions: *d* = base rate of range expansion; *e* = rate of range contraction; *j* = per-event weight of founder-event speciation at cladogenesis (9).

**Table S2.** Nested model test results comparing log-likelihoods (LnL) between nested models from BioGeoBEARS (7). Table shows results from testing for significant model improvement when applying the *j* parameter to each of the biogeographic models.

| Alt. model | Null | LnL alt. | LnL null | DF | D stat. | *p*-value |
| --- | --- | --- | --- | --- | --- | --- |
| **DEC+J** | **DEC** | **-303.1** | **-305.6** | **1** | **4.96** | **0.026** |
| **DIVALIKE+J** | **DIVALIKE** | -361.5 | -361.5 | 1 | -0.0076 | 1 |
| **BAYAREALIKE+J** | **BARYAREALIKE** | -308.6 | -321.8 | 1 | 26.28 | 3.00E-07 |

**Note:** Significant model improvement was inferred based on a *p*-value significance cutoff of >0.05. The *j* parameter allows for founder events within clades (9).

**Table S3.** Comparing morphology-based taxonomic treatments of subgeneric groups of *Asparagus* in southern Africa with species tree clade assignments from this study.

| Current study | Baker (1896) | Jessop (1966) | Obermeyer & Immelman (1992) |
| --- | --- | --- | --- |
| Clade | Section | Section | Series |
| **Africani** | Declinati (1)<br>Umbellati (2)<br>Thunbergiani (1)<br>Africani (1) | Africani (2)<br>Racemosi (1) | Africani (3)<br>Retrofracti (4) |
| **Asparagus** | - | - | - |
| **Exuviali** | Declinati (2) | Exuviali (1) | Exuviali (1) |
| **Macowanii** | Declinati (1) | Africani (2) | Retrofracti (1)<br>Globosi (1) |
| **Myrsiphyllum** | Declinati (1)<br>Striati (1)<br>Myrsiphylli (5) | Africani (1)<br>Racemosi (1)<br>Scandentes (1)<br>Crispi (1)<br>Myrsiphyllum (2) | Myrsiphyllum<sup>a</sup> (11) |
| **Nutlet** | - | Capenses (1) | Suaveolens (3) |
| **Racemose** | Thunbergiani (1)<br>Racemosi (2)<br>Falcati (5) | Racemosi (13) | Africani (1)<br>Exuviali (1)<br>Racemosi (6)<br>Protasparagus (14)<br>Globosi (4) |
| **Retrofracti** | Thunbergiani (3)<br>Africani (2) | Africani (5) | Africani (1)<br>Retrofracti (3)<br>Globosi (3) |
| **Scandens** | Striati (1) | Scandentes (1) | Penduli (1)<br>Myrsiphyllum<sup>a</sup> (1) |
| **Setaceus** | Declinati (2) | Africani (2) | Penduli (2)<br>Retrofracti (1)<br>Globosi (2) |
| **Suaveolens** | Capenses (5) | Capenses (3) | Suaveolens (9) |
| **Sympodioidi** | Umbellati (1)<br>Striati (2) | Striati (2) | Sympodioidi (4) |

**Note:** Parenthesized numbers represent the number of species from each respective, previously described, subgeneric group that were placed in corresponding clades from this study. Baker (10) proposed that all diversity of southern African lineages of *Asparagus* could fit into 9 total sections, whereas Jessop (11) proposed 8 sections and Obermeyer & Immelman (2) proposed 9 series for such.
<sup>a</sup> Represents genus *Myrsiphyllum* Willd.

